# Transferable Transcriptional Topic Modeling Traces Medulloblastoma Subtypes to Distinct Cerebellar Developmental States

**DOI:** 10.1101/2025.11.12.687706

**Authors:** Ashmitha Rajendran, Parthiv Haldipur, Sonali Arora, Kaustubh Grama, Saanvi S. Subramanian, Leyre Merino-Galan, David Johnson, Kimberly A. Aldinger, Jay Shendure, Kathleen J. Millen, John H. Gennari, Siobhan S. Pattwell

**Affiliations:** Department of Biomedical Informatics and Medical Education, University of Washington School of Medicine, Seattle, WA; Ben Towne Center for Childhood Cancer and Blood Disorders Research, Seattle Children’s Research Institute, Seattle, WA; Human Biology Division, Fred Hutchinson Cancer Center, Seattle, WA; Wallace H. Coulter Department of Biomedical Engineering, Georgia Institute of Technology, Atlanta, GA; ⁵Division of Biology and Biological Engineering, California Institute of Technology, Pasadena, CA; Norcliffe Foundation Center for Integrative Brain Research, Seattle Children’s Research Institute, Seattle, WA, USA; Department of Genome Sciences, University of Washington, Seattle, WA, USA; Brotman Baty Institute for Precision Medicine, Seattle, WA, USA; Seattle Hub for Synthetic Biology, Seattle, WA, USA; Howard Hughes Medical Institute, Seattle, WA, USA; Department of Pediatrics, University of Washington, Seattle, WA, USA; Department of Neurology, University of Washington, Seattle, WA, USA

**Keywords:** Topic modeling, single-cell RNA sequencing, cerebellar development, medulloblastoma, SHH subgroup, cell type annotation, pediatric brain cancer, natural language processing

## Abstract

Single-cell transcriptomics transformed our understanding of cellular heterogeneity, yet cross-dataset comparison remains fundamentally limited by batch effects, technology differences, and inconsistent annotation frameworks. These challenges have impeded efforts to connect developmental programs with disease states which creates a critical gap for understanding pediatric cancers with suspected developmental origins. Here, we demonstrate that topic modeling, an unsupervised natural language processing technique, overcomes these barriers by learning transferable transcriptional genetic signatures that generalize across datasets, technologies, and biological contexts without requiring data integration. Applying this framework to over one million fetal cerebellar nuclei, we identify seven distinct topics that capture the developmental spectrum of rhombic lip progenitors through external granule layer (EGL) differentiation, including transitional states missed by conventional clustering. These topics transfer successfully across sequencing technologies (SPLiT-seq to sci-RNA-seq3), developmental timepoints, species (human to mouse), and from single-cell to bulk RNA sequencing of 876 medulloblastoma tumors. We reveal that Sonic hedgehog (SHH) medulloblastoma subtypes retain these topics, corresponding to distinct stages of EGL development. Our findings highlight that developmental timing at tumor initiation fundamentally shapes tumor biology, with immediate implications for subtype-specific therapeutic strategies in pediatric brain cancer. Our workflow establishes topic modeling as a scalable solution for mining expanding genomic atlases with broad applications across different datasets, technologies, and biological contexts.

## INTRODUCTION

Single-cell RNA sequencing (scRNA-seq) has revolutionized the study of cellular heterogeneity in brain development and disease ^1–5^. However, a critical challenge of this method that persists is inconsistent cell type annotations across datasets, which can undermine comparative analysis and translational applications. This gap continues to grow as datasets continuously expand across laboratories, technologies, and biological contexts but our ability to systematically compare and integrate findings has not kept pace. Though powerful statistical and machine learning approaches have been proposed, a recent perspectives of causal machine learning for single cell genomics highlighted several open problems in the field^6^. First, there is a lack of generalization of models to novel experimental conditions; second, there till exists the complexity of interpreting learned models (the “black box”); third, there still is a difficulty of learning cell dynamics instead of one consistent state. Additionally, with the expansion of data, batch correction and data integration methods, while valuable, require careful parameter tuning, can remove genuine biological signals, and become computationally prohibitive^6–9^.

The limitations listed above are consequential for understanding biological problems that require integration of different contexts. For instance, for pediatric brain cancers where origins are strongly tied to developmental cellular dynamics and hierarchies. This precise mapping of tumor-developmental relationships can have downstream effects on therapeutic strategies. The cerebellum is an ideal system for developing such methods as it is the known origin for several pediatric brain tumors and has defined developmental stages. The cerebellar rhombic lip (RL) orchestrates neurogenesis through precisely timed proliferation and differentiation waves. This process over half of all neurons in the adult human brain^10,11^. Beginning around post-conception week (PCW) 8 in humans, specific progenitors migrate from the RL to form the external granule layer (EGL). The EGL is a bilayered germinal zone where granule neuron progenitors (GNPs) undergo massive Sonic hedgehog (SHH)-driven expansion in the outer EGL before transitioning to the inner EGL for differentiation and eventual migration to the internal granule layer^11–13^. This developmental story, though meticulously coordinated, is transient. Its ephemeral existence proves challenging to capture with conventional clustering approaches. These methods are powerful for more prominent cell types but overlook small populations undergoing rapid state transitions that get buried in umbrella clusters. Dysregulation of these EGL developmental processes, particularly aberrant EGL proliferation, can directly contribute to post-natal pathogenesis, specifically medulloblastoma^14–17^.

Medulloblastoma is the most common malignant pediatric brain tumor that arises from these cerebellar progenitor populations. This cancer has four major molecularly defined subgroups: WNT, SHH, Group 3, and Group 4^14^. Seminal work has established that impaired differentiation of specific neural progenitors underlies pediatric brain cancers^18^. By projecting bulk tumor transcriptomes onto developmental single cell atlases, Vladoiu et al. demonstrated that medulloblastoma subgroups match to specific developmental cellular lineages. Here, they revealed highly defined WNT medulloblastomas map to rhombic lip derived mossy fiber neurons. Similarly, from another pivotal integrative multi-omic work from Northcott et al^17^, Group 3 and Group 4 medulloblastoma tumors mapped to photoreceptor and unipolar brush cell expansions from the RL. Another complementary work from Hendrikse et al^19^ employed unique approaches to map tumor cells of origin through bulk transcriptomics projected onto single cell developmental atlases, DNA methylation, and enhancer landscape analysis and further validated medulloblastoma tumor subgroup origins.

While these approaches established the developmental origins of medulloblastoma subgroups, there are several unaddressed challenges. First, projection-based methods rely on pre-defined developmental signatures which miss novel information or skew our understanding of existing sell types. Second, these require integration across disparate datasets with different sequencing technologies, processing pipelines, and annotation schemes. These integration and data harmonization methods invariably lose data and can oversimplify the existing biology. Finally, the developmental populations that are the most relevant to tumor origins are small, transient, and poorly represented in most of these atlases which prevents more robust and detailed mapping. These methodological constrains highlight the need for analytical frameworks that can extract transferable biological information without requiring direct data integration.

We demonstrate here that topic modeling^20^, a natural language processing approach, can treat cells as “documents” and genes as “words”^21^ in order to identify conserved transcriptional programs across diverse datasets without requiring data integration or shared processing pipelines (analogy demonstrated in Figure S1 and workflow demonstrated in Figure S2). Topics from one dataset can be directly applied to others, enabling connections between developmental programs and cancer. While different studies might use varying labels for similar populations, topic modeling reveals shared or distinct transcriptional programs beneath annotation disagreements. Ultimately, we show here that this topic modeling framework identifies biologically meaningful gene expression patterns that persist across technologies, timepoints, and between healthy and malignant tissues.

As proof of principle, we analyzed midgestational fetal cerebellar landmark datasets. The Aldinger et al. dataset^1^ (subset to 15,556 nuclei from post-conception week 17) serves as our source for establishing cerebellar-specific transcriptional programs, benefiting from expert annotation. Post-conception week 17 represents an ideal developmental timepoint for identifying EGL-associated transcriptional programs, as the EGL is well-established and actively expanding at this stage, with robust populations of both proliferating oEGL and differentiating iEGL granule neurons. The Cao et al. dataset^2^ (subset to 119,954 nuclei from weeks 12.5) serves as our validation set for testing topic conservation across different experimental approaches. Week 12.5 was selected as it contained the largest number of nuclei in the Cao dataset and represents an earlier developmental stage where we would expect distinct distributions of developmental topics. This distribution would be focused specifically with greater enrichment of RL-associated programs and proportionally less EGL activity compared to the later week 17 timepoint. This temporal comparison allows us to capture the developmental transition from RL-dominant to EGL-dominant neurogenesis. For disease context, we use the Arora et al. dataset which encompasses medulloblastoma bulk RNA sequencing^22^ across multiple subgroups, examining how developmental programs persist or alter in malignancy.

To further validate topic conservation, we applied our topics to a mouse developmental atlas from Qiu et al.^23^ and examined expression patterns of top topic genes using mouse in situ hybridization. These analyses confirmed EGL localization and validated the genetic signatures proposed here as capturing transitional cell states within the EGL This also demonstrates the ability of our pipeline to combine and analyze data across three labs, different genomic data, and collection technologies.

Our analysis shows several advances. First, topic modeling identifies transitional states and rare cell types, particularly developing human EGL progressions which includes oEGL to iEGL transitions and migration of differentiating granule neurons. These specific transitions are missed by conventional clustering approaches. This captures the spectrum of GNP proliferation, cell cycle exit, and early differentiation within the EGL structure. Second, topic transfer using gene set variation analysis (GSVA) successfully identifies conserved cell types and developmental processes across independent developmental datasets from different labs and technologies. The transferability demonstrates robust detection of EGL associated genetic signatures across developmental timepoints. Third, applying cerebellar developmental topics to medulloblastoma bulk RNA sequencing confirms the GNP and EGL origins of SHH tumors and reveals age-specific molecular signatures within SHH subtypes. This disease application demonstrates the transferability of topics across data types and biological contexts. Notably, the EGL-derived topics allow us to distinguish which stage of granule neuron development, such as proliferative oEGL versus differentiating iEGL, is most represented in different SHH medulloblastoma subtypes. A major contribution to the field here is the generalizability of this pipeline, which enables systematic mapping of EGL developmental programs onto disease states. These findings improve our understanding of cerebellar neurodevelopment, particularly the molecular architecture of the EGL, and guide downstream characterization of tumor substructure that can inform and alter treatment strategies.

## RESULTS

### Topic modeling identifies conserved transcriptional programs across cerebellar development datasets

Our computational framework transfers developmental transcriptional signatures (topics) across datasets and biological contexts without requiring data integration. Figure 1 illustrates an overview of the workflow: topics are derived from the source dataset (Aldinger et al., PCW 17), validated in an independent developmental dataset (Cao et al., PCW 12.5), and applied to medulloblastoma bulk RNA-seq to connect developmental programs with disease states.

**Figure 1.**
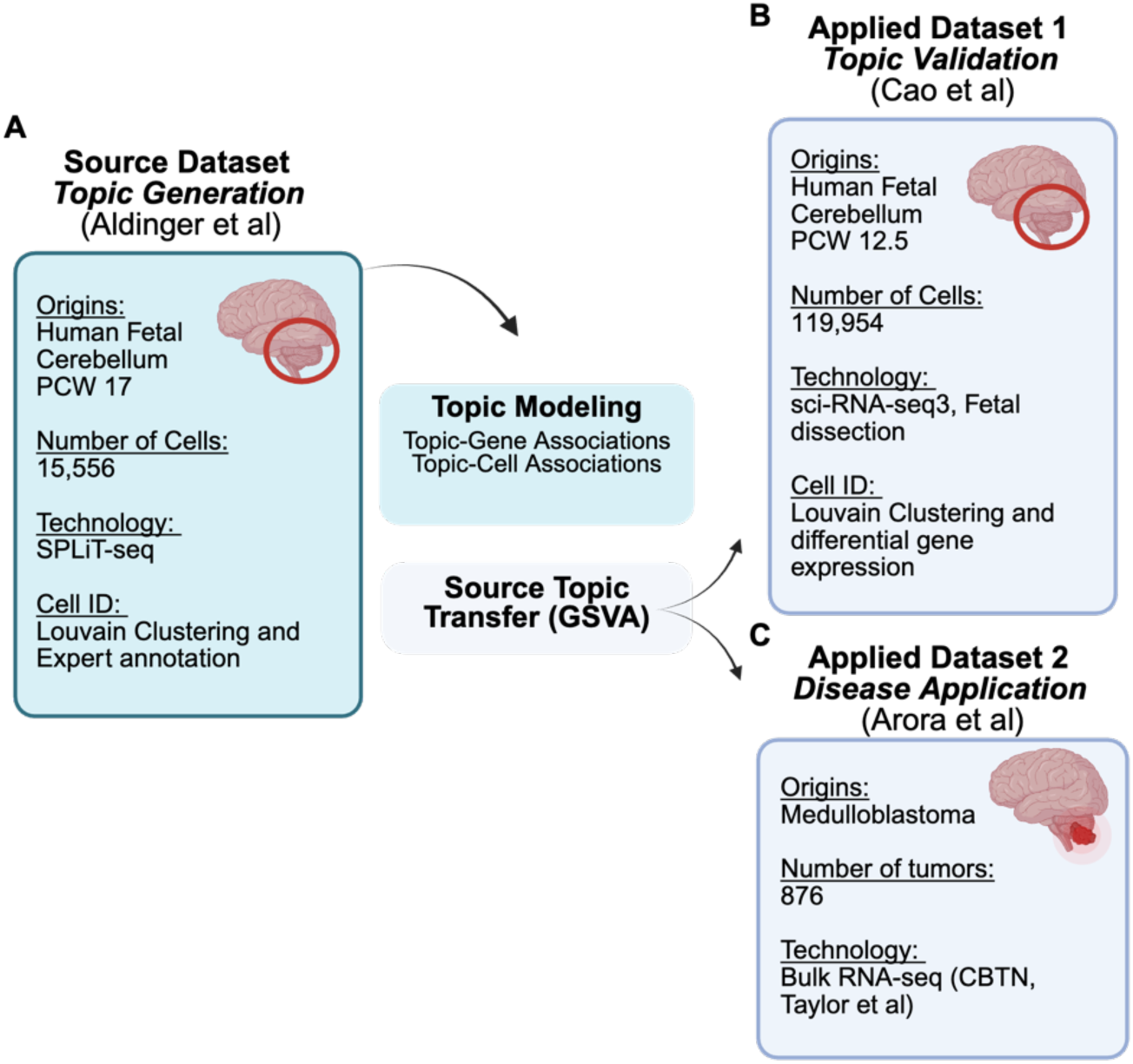
Computational workflow for topic modeling and transfer with from Source dataset to two Applied datasets. (A) Source Dataset (Aldinger et al., subset of 15,556 nuclei from PCW 17 of human cerebellar development) (B) Applied Dataset 1 (Cao et al., subset of 119,954 cells from PCW 12.5 of human cerebellar development) (C) Applied Dataset 2 (medulloblastoma bulk RNA sequencing).

Analysis of the Aldinger et al. dataset (subset to 15,556 nuclei from post-conception week 17) (Figure 1A) using topic modeling identified 65 topics. Our method included the use of iterative modeling to select the optimal number of topics based on perplexity (Table S1 and see Methods). Perplexity is a statistical measurement that quantifies how well a model predicts on held-out data. By selecting a minimum plateau in perplexity across iterations allows for selecting a model with less redundancy while maintaining complexity. Initially, these topics are given arbitrary numeric identifiers that we then renamed based on biological function.

Then, we transferred the source topics to the independent Cao et al. dataset (subset to 119,954 nuclei from week 12.5) using GSVA (Figure 1B) of the top 50 genes per topic from the source dataset (Table S3), these topics showed conservation despite differences in technology (SPLiT-seq vs sci-RNA-seq3) and developmental timepoints, validating their biological relevance. These topics were then tested on Applied dataset 2 in a medulloblastoma application to verify that the same topics related to GNPs and the EGL are also identifiable in medulloblastoma bulk RNA sequencing (Figure 1C). This dataset is a reference landscape for human brain disease of which we subset for medulloblastoma (n=876 tumors).

To specifically identify topics that may represent the EGL, we (1) identified topics with the presence of known EGL marker genes and pathways among their top-ranked gene-topic associations, (2) compared topic-gene associations with the top differentially expressed genes (DEGs) from the EGL through hierarchical clustering analysis, and (3) examined GSVA-based UMAP localization to granule neuron, granule cell progenitor, and rhombic lip cell type clusters in the source dataset with existing expert annotations as well as follow up expert validation. The selected topics were then validated across independent datasets using GSVA-based topic transfer and through spatial validation via in situ hybridization of top marker genes from each topic in mouse cerebellar tissue.

### Transition Cell States in the External Granule Layer (EGL)

The resulting 65 topics derived from the source dataset included a subset that captured the EGL which we identified through combinations of EGL-based genetic markers and RL-derived lineages within the EGL (Figure 2), identified through the presence of known RL marker genes and EGL genes. The cerebellar EGL is organized into distinct zones from the pial surface to the internal granule layer (IGL). The outer EGL contains mitotic and proliferative cells, including early transient Rhombic Lip-derived (RL) and locally proliferating progenitors (Topic 22) and intermediate progenitors (Topic 2). The transition zone harbors differentiating neuronal cells. The inner EGL consists of post-mitotic and pre-migratory neurons, including differentiating cells (Topic 49), early differentiated granule neurons (GNs) (Topic 46), and post-mitotic cells (Topic 61). Mature neurons migrate from the inner EGL toward the IGL, with transitioning mature GNs represented by Topic 25. Glial progenitors (Topic 27) span multiple zones throughout this developmental trajectory.

**Figure 2.**
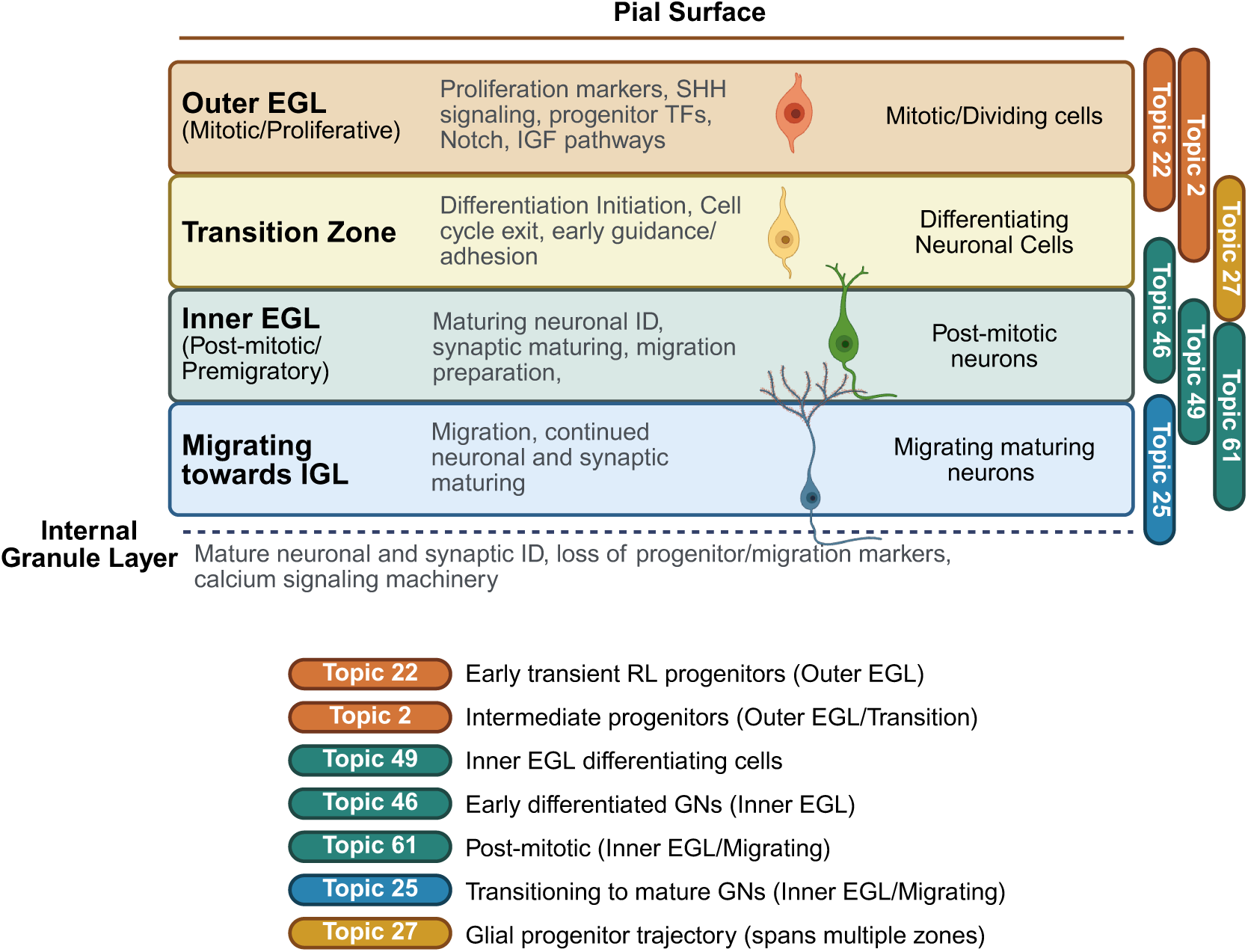
Spatial organization of neurogenic and gliogenic cell populations across the external granule layer during cerebellar development. Schematic representation of cerebellar EGL structure showing the distribution of seven EGL-associated topics (Topics 2, 22, 25, 27, 46, 49, 61) across developmental zones from outer EGL to internal granule layer. Topics are color-coded and labeled according to their spatial localization patterns.

These topics were selected from the pool of 65 topics through three complementary approaches. First, these topics were identified through the presence of known EGL marker genes and pathways among their top-ranked gene-topic associations, including proliferation markers *MKI67* and *CENPF*, SHH pathway component *GLI2*, and the human-specific *ARHGAP11B* associated with progenitor expansion, alongside EGL differentiation markers for the neuronal lineage topics. Second, hierarchical clustering analysis comparing topic-gene associations with the top differentially expressed genes (DEGs) from the EGL identified in the Aldinger et al. study showed that Topics 49, 22, 2, 27, and 46 showed the strongest selective enrichment for these EGL-specific markers (Figure S3). Third, we examined GSVA-based UMAP localization to granule neuron, granule cell progenitor, and rhombic lip cell type clusters (Figure 3A) in the source dataset. The resulting topics begin to reveal previously unrecognizable substructure within the EGL developmental continuum. They capture cellular heterogeneity that conventional clustering methods struggle to resolve without extensive expert knowledge and manual curation.

**Figure 3.**
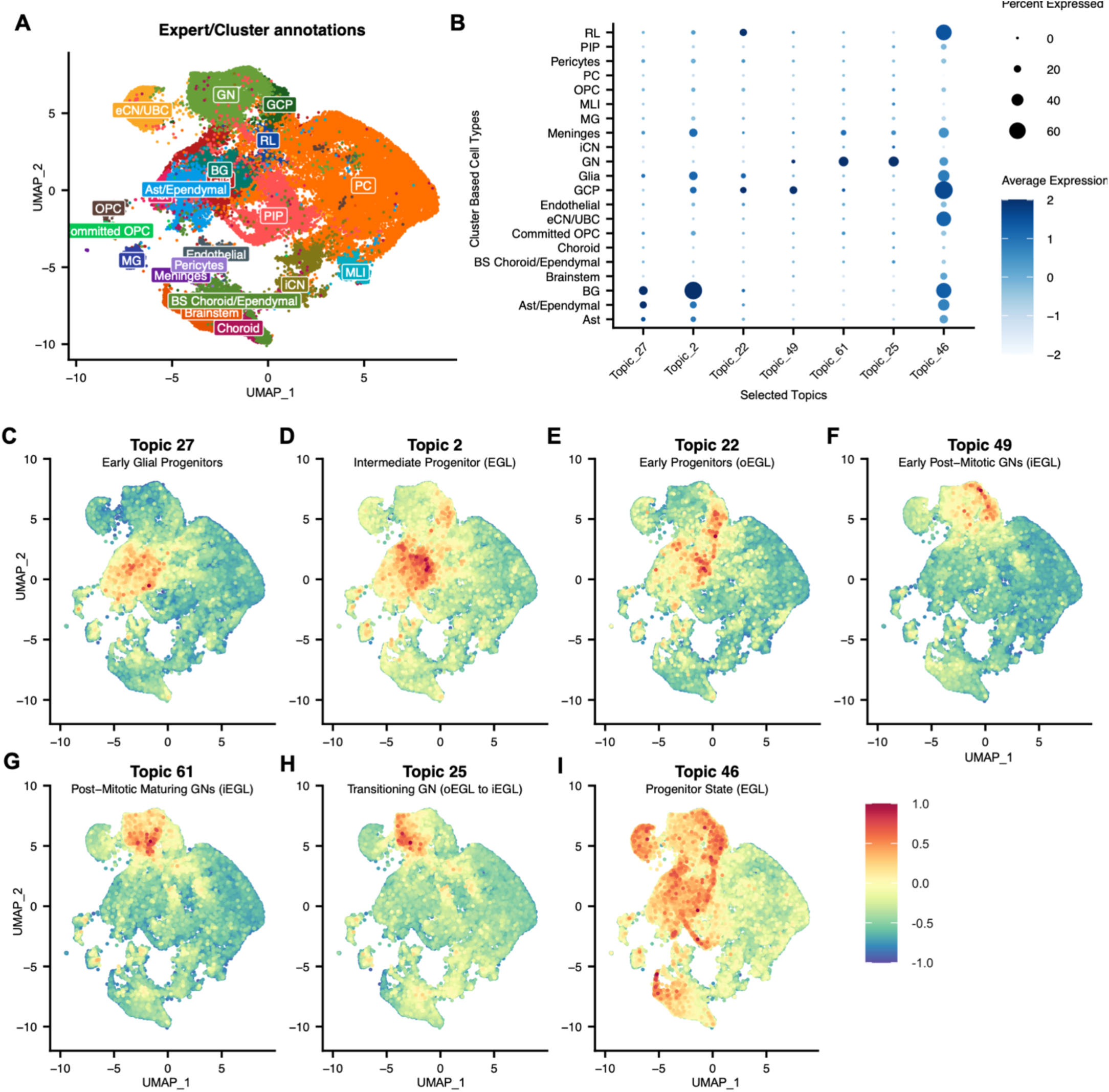
Topic modeling reveals distinct transcriptional programs defining rhombic lip progenitor lineage trajectories in human cerebellar development. (A) UMAP visualization of 15,556 nuclei from post-conception week 17 (Aldinger et al. source dataset)^1^ colored by original cell type annotations, including rhombic lip (RL), granule cell precursors (GCP), granule neurons (GN), Purkinje cells (PC), molecular layer interneurons (MLI), cerebellar nuclei neurons (iCN/eCN), oligodendrocyte precursors (OPC), committed OPCs, astrocytes (Ast), Bergmann glia (BG), microglia (MG), and vascular/meningeal populations. (B) Dot plot showing average expression (color intensity) and percentage of cells expressing (dot size) six key topics across major preannotated cerebellar cell types. Topics capture the rhombic lip-to-glial (Topics 27, 2) and rhombic lip-to-neuronal (Topics 22, 49, 61, 25) developmental trajectories. (C-H) UMAP projections of individual topic expression scores from GSVA analysis. Color scale represents scaled GSVA scores. (C) Topic 27 (Early Glial Progenitors) (D) Topic 2 (Intermediate Progenitor (EGL)) (E) Topic 22 (Early Progenitors (oEGL)) (F) Topic 49 (Early Post-Mitotic GNs (iEGL)) (G) Topic 61 (Post-Mitotic Maturing GNs (iEGL)) (H) Topic 25 (Transitioning GN (oEGL to iEGL)

### Early Proliferative Rhombic Lip Progenitors in the Outer EGL (Topic 22)

Topic 22 represents an early, transient rhombic lip-derived early proliferating progenitor population genetic signature marked by proliferation genes *MKI67*, *CENPF*, and *PARD3* (centrosome marker) indicating active cell division. This topic also contains *GLI2* (essential for SHH signaling). The presence of *PARD3*, a subpolarity gene, is consistent with cells undergoing active spatial reorganization and is critical for germinal zone exit (Famulski et al., 2010; Singh et al., 2016). Additional proliferation and cell division genes includei*CDK6* (cell cycle regulator), *EZH2* (chromatin remodeling), *MPPED2*, *DIAPH3* (cytoskeleton regulation), *STARD13*, *KNTC1* (kinetochore function), and *CIT* (cytokinesis). This population may be associated with cerebellar foliation processes, as evidenced by the presence of *ARHGAP11B*, a human-specific gene that has been linked to increased cerebral gyration and cerebellar foliation^24–26^. In addition, Topic 22 has its strongest expression in the rhombic lip (RL) with very minimal expression across other cell types (Figure 3B). consistent with its identity as an early progenitor signature. The topic-cell association values projected using UMAP (Figure 3E) shows Topic 22 is sharply localized to a small discrete population within the RL cluster (upper left region), with virtually no expression in the large GN/GNP cluster. This type of pattern reflects the transient and spatially restricted nature of this progenitor state at this developmental timepoint. The exact temporal relationship between Topic 22 and Topic 2 cannot be definitively ordered, as both represent early transient states.

### Intermediate Progenitor State within EGL Transition (Topic 2)

Topic 2 represents an intermediate progenitor state starting to transition from the outer EGL to differentiated populations, characterized by expression of *SOX2, MEIS1, PAX3*, and *NFIA*. *MEIS1* is a transcription factor that promotes BMP signaling and *ATOH1* degradation, driving cell cycle exit and differentiation^11,27^. This topic contains SHH pathway components (*PTCH1*, *GLI2*) essential for glia survival^11,28,29^. The co-expression of *GRIA4* (astrocyte-associated), *SOX2-OT* (glial marker), and *NFIA* (downstream of *PAX3* in oligodendrocyte precursor specification) suggests this captures progenitors at a critical lineage decision point. The proliferation signature (*MKI67, CENPF*) combined with *GLI2* expression without *LMX1A/ATOH1* confirms its differentiating EGL progenitor identity. Like Topic 22, this population contains *PARD3* expression consistent with its transitional nature and may be present during active cerebellar foliation associated with *ARHGAP11B* expression. Additionally, Topic 2 has the highest expression in Bergmann glia (BG) and astrocytes (Ast/Ependymal, Ast), with moderate expression in glia and GNP, suggesting a glial-progenitor intermediate state (Figure 3B). Through scaled topic-cell expression patterns (Figure 3D), Topic 2 is enriched in both the glial clusterand portions of the GN/GNP cluster, with particularly strong expression at the boundary between these populations, consistent with cells at a progenitor lineage decision point bridging neuronal and glial fates.

### Glial Progenitors diverging from Neuronal EGL Lineage (Topic 27)

Topic 27 represents an early glial progenitor trajectory located in the outer rhombic lip, enriched in radial and Bergmann glia annotated and related progenitor populations. This topic is identifiable with markers *PTN, SMOC1, TNC,* and *ETV5* (apical, basal radial and Bergmann glial markers) alongside *SLC1A3* (definitive glial transporter) and *GDF10* (glial marker). The presence of *PAX3* and downstream effector *NFIA* links this population to oligodendrocyte precursor specification^1^. Additionally when compared to existing cell type annotatoins (Figure 3A-B), Topic 27 showed highest expression in Bergmann glia (BG), astrocytes (Ast, Ast/Ependymal), and radial glia, with some expression in RL, suggesting a glial progenitor identity. This is reflected in the scaled GSVA values (Figure 3C) where Topic 27 is strongly enriched in the large glial/astrocyte cluster, with minimal expression in the GN/GCP cluster, demonstrating clear spatial segregation consistent with commitment to the glial rather than neuronal fate. This topic likely represents early glial progenitors distinct from the neuronal lineage trajectories captured by the other topics.

### Early Post-Mitotic Cells in Inner EGL initiating Differentiation (Topic 49)

Topic 49 contains a geneset that represents a population of recently differentiated cells within the inner EGL. These cells are characterized by expression of NEUN *(RBFOX3), RELN, PAX6*, and *ROR*, consistent with their post-mitotic identity. Notably, *RELN* expression shows enrichment in the outer relative to the inner EGL, a pattern confirmed in in situ hybridization of mouse brain at P4 from the Allen Cell Atlas (Figure S6C). The very low gene-topic association of *ATOH1* suggests these are transitioning cells that have moved beyond the early progenitor stage^30^. The presence of *NEUROD1*, which is initially expressed during neuronal differentiation, indicates that Topic 49 likely represents cells upstream of *NEUROD1* expression in the differentiation trajectory. This population also expresses synaptic adhesion molecules (*LRRTM4, NLGN1, NLGN4X*) and guidance cues (*ROBO2, DSCAM*), suggesting these are migrating granule neuron progenitors beginning synaptic organization^1^. The scaled GSVA values (Figure 3F) reveals that topic 49 is localized to specific regions within the granule neuron cluster from the cluster annotations. This is consistent with cells that would be at an early post-mitotic stage of differentiation.

### Inner EGL Post-Mitotic Neurons with developing Synaptic Connectivity (Topic 61)

Topic 61 represents a distinct inner EGL genetic signature characterized by *RBFOX3* (NEUN) expression, which marks post-mitotic neurons and definitively excludes these cells from the outer EGL compartment (Lee et al., 2020). This population expresses glutamate receptors (*GRIK2, GRIA4*), synaptic proteins (*RELN*), and calcium signaling components (*RYR3, PRKCB*), indicating these cells are establishing functional connectivity while still undergoing maturation. This population appears to represent a slightly later stage in the differentiation process compared to Topic 49, representing another transition state within the inner EGL as granule neuron progenitors commit to their neuronal fate.These populations of cells that have high expression of this topic may represent novel “states” of EGL cells that have not been previously well-characterized. Topic 61 is also enriched in intermediate regions of the granule neuron cluster (Figure 3G), that is consistent with cells that have progressed beyond early differentiatoin but have not completed maturation.

### Maturing Neurons Transitioning from inner EGL towards Internal Granule Layer (Topic 25)

Topic 25 represents a more differentiated neuronal state among these populations, characterized by *ROBO1* (definitive EGL marker), mature synaptic proteins (*UNC13C*), and neuronal specification factors (*RBFOX1*, *MEG3*). This population also expresses DSCAM (granule neuron marker), *RELN* (EGL attractant), and *GRIK2* (Lee et al., 2020). When compared to the existing annotations of the cells (Figure 3B), Topic 25 showed strong association with granule neurons (GN) and moderate expression in eCN/UBC, suggesting a mature neuronal signature. From UMAP visualization of the scaled GSVA values (Figure 3H), Topic 25 is enriched throughout much of the GN cluster with a relatively uniform distribution, extending into some adjacent populations, consistent with cells that have completed migration and are integrating into cerebellar circuits. These cells may represent an intermediate state between the inner EGL and fully differentiated granule neurons that will eventually migrate to form the internal granule layer^1^.

### Progenitor State with sustained plasticity in EGL (Topic 46)

Topic 46 represents a slightly distinct population marked by *MEIS1* (which promotes cell cycle exit and differentiation^27^), *PDE1A* (expressed throughout granule cell development), and *DACH1* which are all markers of EGL identity. However, this population also expresses neural developmental transcription factors including *ERBB4, NFIA, NFIB, TCF4*, and *CHD7* (*CHD7* promotes clonal expansion of granule neuron progenitors^11,31^). *ERBB4* is found in the inner EGL and in regions distinct from Purkinje cells, suggesting these may be among the earliest born granule neurons. The expression of *ZEB1* is particularly notable, as this factor controls neuron differentiation and germinal zone exit through regulation of polarity genes including *PARD3* and *PARD6*^32^. *ZEB1* is downregulated as GNPs exit the EGL, and its persistent expression delays morphological maturation and migration to the IGL. Importantly, *ZEB1* expression is elevated in SHH medulloblastoma and has been identified as a subgroup-specific marker of prognosis and potential therapeutic target^33^. This topic also contains synaptic adhesion molecules (*LRRTM4, NLGN1, NLGN4X*) and the CSMD family of cell surface glycoproteins (*CSMD1, CSMD2, CSMD3*), along with additional neural development markers including *LHFPL3*, *NTNG1* (axon guidance), *AGBL4*, and *STK32B*.

When compared to preexisting cell type annotations, Topic 46 had strongest expression in eCN/UBC, GCP, and BG, with notable expression also in RL and glia, indicating a broad expression pattern across multiple cell types (Figure 3B) though strongly in the RL. Additionally, Topic 46 (Figure 3I) has diffuse enrichment spanning the RL region, portions of the GN/GCP cluster, and extending into glial populations, confirming its expression across multiple cell clusters, consistent with a rare or multipotent progenitor state. These findings represent some of the first clear distinctions of granule neuron proliferation and differentiation states, particularly the transition between outer and inner EGL compartments.

### Cross-dataset validation confirms robust conservation of developmental programs

To assess whether topics generated using the Source data (Figure 1A) can be used to identify similar developmental programs in a different dataset (Figure 1B), regardless of how that dataset was processed, we applied source topics to applied dataset 1 (Cao et al.)^2^. Applied dataset 1 is another human developmental cerebellar dataset using sci-RNA-seq3 where we used subset for PCW 12.5 (Figure 4A). Here, we focus on the topics above that overlap with the EGL lineages: Topics 27, 2, 22, 49, 61, and 25. These topics showed consistent enrichment patterns (Figure 4B) despite differences in sequencing technology (SPLiT-seq vs sci-RNA-seq3) and specimen ages.

**Figure 4.**
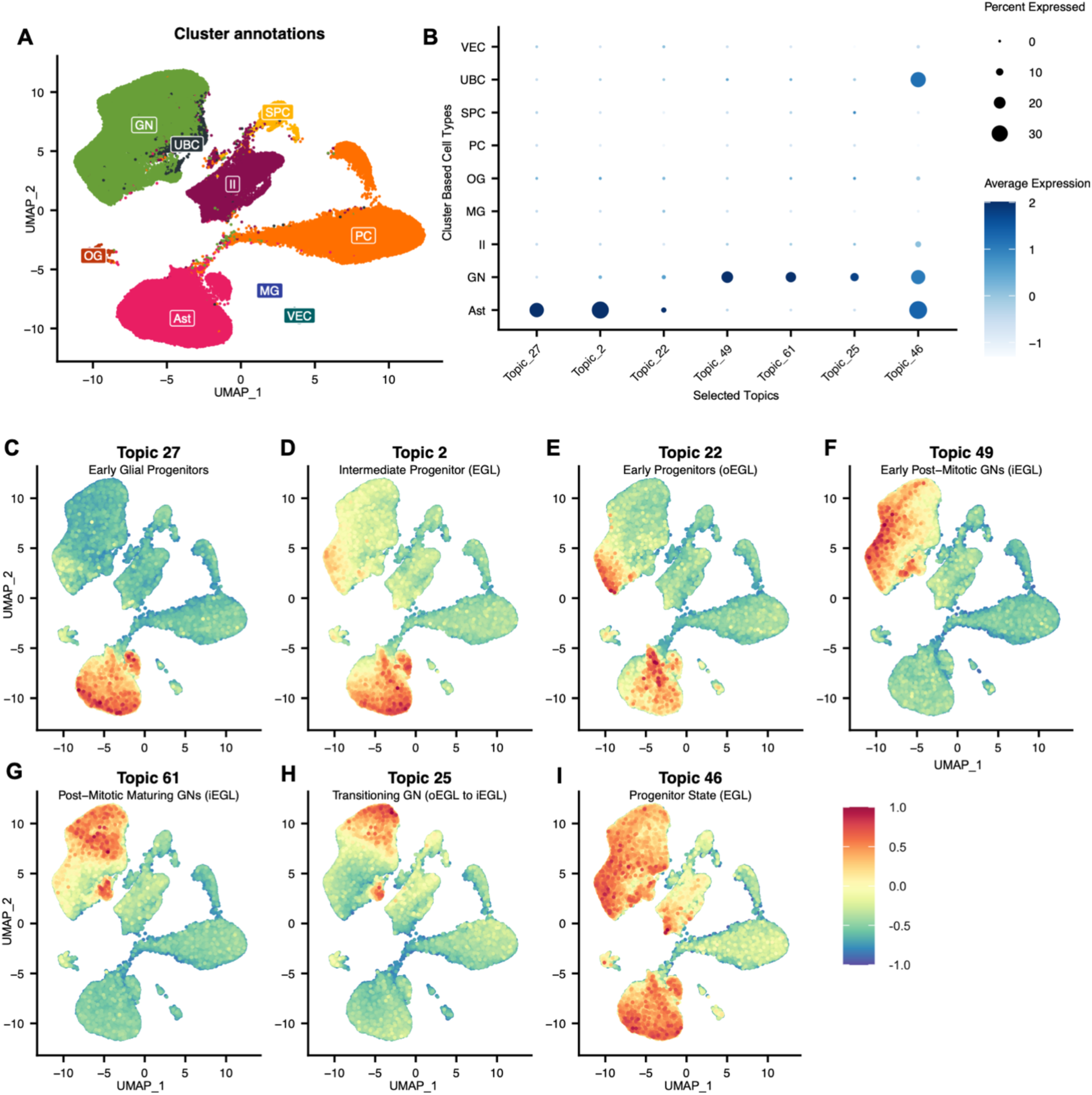
Cross-dataset validation demonstrates robust conservation of rhombic lip lineage topics from source dataset in an independent human developmental cerebellar dataset (applied data). (A) UMAP visualization of 15,556 nuclei cells from post-conception week 12.5 (Cao et al. applied dataset)^2^ colored by original cell type annotations, including granule neurons (GN), unipolar brush cells (UBC), astrocytes (Ast), Purkinje cells (PC), oligodendrocytes (OG), microglia (MG), molecular layer interneurons (II), vascular endothelial cells (VEC), and SLC24A4+/PEX5L+ cells (SPC). (B) Dot plot showing average topic expression (color intensity) and percentage of cells expressing (dot size) for the same six rhombic lip lineage topics identified in the source dataset, now applied to this independent dataset via GSVA transfer. (C-H) UMAP projections of individual topic expression scores following GSVA-based topic transfer. Color scale represents scaled GSVA scores. (C) Topic 27 (Early Glial Progenitors) (D) Topic 2 (Intermediate Progenitor (EGL)) (E) Topic 22 (Early Progenitors (oEGL)) (F) Topic 49 (Early Post-Mitotic GNs (iEGL)) (G) Topic 61 (Post-Mitotic Maturing GNs (iEGL)) (H) Topic 25 (Transitioning GN (oEGL to iEGL)

Topic 27 (Figure 4C) and Topic 2 (Figure 4D) showed enrichment patterns consistent with glial progenitors and intermediate EGL states, respectively. On average, they both congregated in the cluster previously identified broadly as ‘astrocytes’ (Figure 4B). However, they both identify substructure within this cluster and show overlap with part of the granule neuron cluster at different intensities (Figure 4C-D).

Topic 22, which we identified as marking early progenitors in the oEGL, identified a distinct population in the Cao dataset that had not been annotated as such in the original study. This population is located between the clusters previously identified as astrocytes and granule neurons which is the expected region where rhombic lip derived early progenitors would exist (Figure 4E).

Topics 49 (Figure 4F), 61 (Figure 4G), and 25 (Figure 4H), which capture the progression from early neuronal differentiation into maturation, reveals previously unrecognized substructure within the granule neuron cluster in the Cao dataset. Topic 49 consistently marked cells at the periphery of the granule neuron cluster, suggesting these cells represent newly differentiating cells beginning their migration from the EGL. Topic 61 is enriched in the intermediate regions, while Topic 25 is concentrated in the cluster core, indicating the most mature granule neurons. This spatial organization within the UMAP recapitulates the known developmental trajectory, providing orthogonal validation of these topic identities.

### Spatial validation confirms EGL localization of topic marker genes

To validate that our computationally-derived topics capture genuine developmental programs with anatomical correlates in the EGL, we examined the spatial expression patterns of key topic marker genes using in situ hybridization from the Allen Developing Mouse Brain Atlas^34,35^ at postnatal day 4. We selected P4 as it represents a developmental stage when the mouse EGL is well-established and actively producing granule neurons, analogous to the mid-gestation human cerebellar development captured in our source dataset. As a secondary validation, we confirmed that human topics successfully transferred to mouse cerebellar single-cell data (Supplementary Figure S4, using gene conversion Table S4), where they localized to predicted cell types in the developmental mouse atlas (Figure S4A-B), demonstrating cross-species conservation of these transcriptional programs. Each of the topics was projected onto the mouse developmental dataset and scaled GSVA scores begin to find substructure in the broader cell type categories in the CNS (Figure S4C-I).

For spatial validation, we systematically examined nine genes representing core markers across our EGL-associated topics available through the atlas, focusing on their localization to EGL versus non-EGL compartments.

### Spatial validation of Pan-EGL and Transitional Markers

PAX6 is a canonical marker of RL derivatives expressed throughout granule neuron development. PAX6 showed robust, specific expression throughout the EGL with particularly strong signal in both outer and inner zones (Figure 5A), confirming its role as a pan-EGL marker. Computationally, PAX6 showed strong association with Topic 49 (association value: 14,193), which represents early post-mitotic granule neurons in the inner EGL. The strong association to PAX6 validates our topic assignment to the EGL and in conjunction with the other top markers of Topic 49 we can distinguish oEGL proliferative states from iEGL differentiating states.

**Figure 5.**
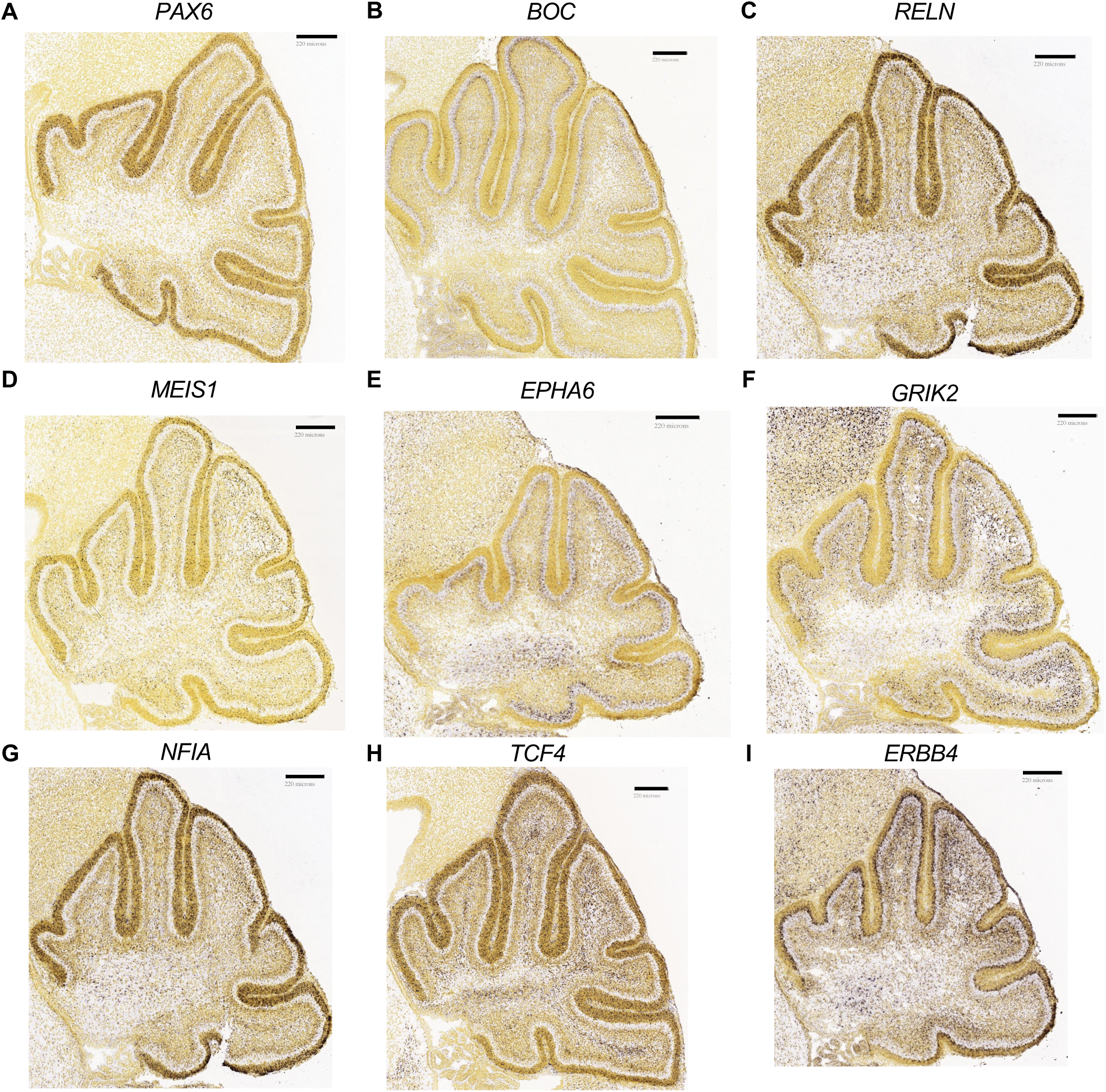
Spatial validation of EGL-associated topic marker genes in developing mouse cerebellum at postnatal day 4. In situ hybridization images from the Allen Developing Mouse Brain Atlas showing spatial localization of genes computationally identified as markers of EGL-associated topics. Gene-topic associations are based on association values from Table S2. Scale bars = 220 μm. (A) PAX6 (B) BOC (C) RELN (D) MEIS1 (E) EPHA6 (F) GRIK2 (G) NFIA (H) TCF4 (I) ERBB4

BOC (brother of CDO), a component of the Hedgehog signaling pathway, displayed expression in the inner EGL with some extension into the outer EGL (Figure 5B), consistent with its role in maintaining SHH responsiveness in GNPs. Computationally, BOC showed strongest associations with Topic 2 (17,698) representing intermediate progenitors in the EGL, Topic 27 (15,636) marking glial progenitors, and Topic 22 (12,736) representing early progenitors in the outer EGL, supporting its expression across multiple EGL compartments and progenitor lineages.

RELN (Reelin), strongly associated with Topics 61 (50,282), 49 (24,603), and 25 (36,146) representing the progression from inner EGL post-mitotic neurons through mature granule neurons, revealed critical spatial organization within the EGL. RELN showed robust expression throughout the EGL with enrichment in the outer EGL relative to the inner EGL (Figure 5C). This outer-to-inner gradient validates our topic assignments that distinguish oEGL proliferative states from iEGL differentiating states, demonstrating that our topics capture spatial organization within the EGL structure.

MEIS1, strongly associated with Topic 2 (33,190) representing intermediate progenitors and Topic 46 (24,443) marking progenitors with sustained developmental plasticity, exhibited expression throughout the EGL with particular enrichment in differentiating zones (Figure 6D). This spatial pattern is consistent with MEIS1’s known role in promoting cell cycle exit and differentiation, validating Topic 2’s identity as an intermediate progenitor at the transition from proliferation to differentiation.

**Figure 6.**
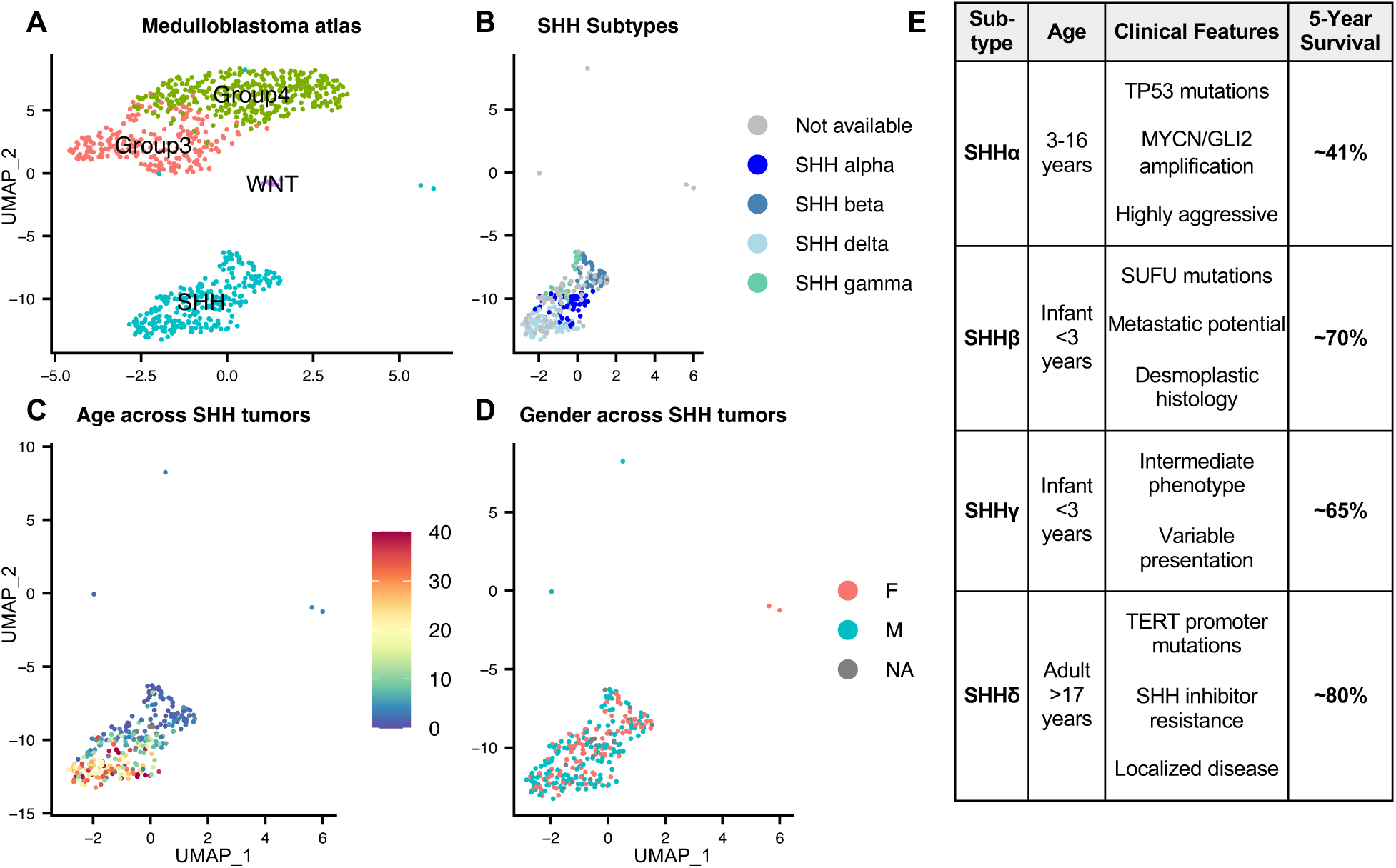
Atlas of medulloblastoma bulk RNA sequencing (Applied dataset 2). (A) UMAP visualization of 876 medulloblastoma tumors colored by molecular subgroup: WNT (n=70), SHH (n=274), Group 3 (n=144), and Group 4 (n=388). (B) UMAP of SHH subgroup subset showing four molecular subtypes (SHHα, SHHβ, SHHγ, SHHδ). (C) UMAP of SHH subgroup subset colored by age distribution across tumors. (D) UMAP of SHH subgroup subset colored by gender distribution across tumors. (E) Clinical characteristics table summarizing age ranges, molecular features, clinical behavior, and 5-year survival rates for each SHH subtype.

### Spatial Validation of Inner EGL and Transitioning Populations

GRIK2 strongly associated with Topics 61 (53,729) and 25 (37,713) represents inner EGL post-mitotic maturing granule neurons and transitioning granule neurons, showed expression in the inner EGL and differentiating granule neurons with signals extending from the iEGL into regions preparing for migration (Figure 5F). The spatial distribution of this synaptic marker, concentrated in the iEGL, confirms that Topics 61 and 25 capture sequential stages of functional maturation as granule neurons exit the germinal zone and establish synaptic connectivity.

EPHA6 associated with Topic 25 (36,018) represents transitioning and maturing granule neurons, displayed expression in the IGL (Figure 5E), consistent with more mature, post-migratory granule neurons that have completed their transition from the EGL. This localization validates Topic 25’s identity as capturing the late stages of granule neuron maturation and migration.

ERBB4, showing strong computational associations with Topics 49 (46,196), 61 (40,835), and 25 (30,986) spanning the inner EGL through transitioning populations, as well as Topic 46 (15,232) marking progenitors with sustained plasticity, displayed robust expression in the inner EGL (Figure 5I). This spatial pattern supports ERBB4’s role in marking granule neurons undergoing differentiation and early maturation within the iEGL compartment, consistent with its presence in multiple topics capturing the iEGL-to-migration transition.

### Spatial validation of Progenitor and Early Differentiation Markers

NFIA is a transcription factor that has a high topic-gene association with Topic 46 (43,944), which represents progenitors with sustained plasticity, Topic 49 (19,015) marking early post-mitotic neurons, and Topic 2 (13,044) representing intermediate progenitors, displayed strong expression throughout the EGL in progenitor zones (Figure 5G). This spatial pattern validates NFIA’s role in maintaining developmental flexibility across multiple progenitor and early differentiation states captured by our topics.

TCF4, a neural developmental transcription factor showing strongest computational association with Topic 46 (42,576) representing progenitors with sustained developmental plasticity, showed expression in progenitor populations within and adjacent to the EGL (Figure 5H). This spatial pattern supports Topic 46’s identity as marking cells that retain developmental flexibility even as they progress through early differentiation stages.

Across all nine genes examined, spatial expression patterns were consistent with predicted topic associations and developmental functions. EGL-enriched genes (PAX6, BOC, RELN, MEIS1, NFIA, TCF4) showed robust localization to the germinal zone, with several displaying layer-specific patterns (outer vs inner EGL) that correspond to the oEGL-to-iEGL progression captured by our topics, specifically Topics 2, 22, 27, and 46). Genes associated with neuronal maturation (GRIK2, EPHA6, ERBB4) showed expression patterns consistent with the inner EGL through post-migratory populations, validating the sequential developmental stages captured by early (Topic 2, 22, 27, 46) through later transitions (Topics 49, 61, and 25).

This spatial validation begins to demonstrate that our topic modeling framework successfully identifies biologically meaningful and anatomically organized developmental programs within the EGL. The concordance between computational topic assignments (based on gene-topic association values from Table S2) and physical tissue localization (from in situ hybridization) provides orthogonal confirmation that extends beyond computational predictions.

### SHH medulloblastoma retains human EGL transition states

To validate our framework’s ability to bridge developmental and cancer contexts, we applied cerebellar developmental topics from the source dataset to medulloblastoma samples from a published bulk RNA sequencing atlas^22^ which includes 876 medulloblastoma tumors (274 labeled as SHH). The UMAP visualization reveals clear separation of the four molecular subgroups, WNT, SHH, Group 3, and Group 4, with SHH forming a distinct cluster (Figure 6A). Within the 274 SHH tumors, the dataset is further divided into four subtypes (SHHα, SHHβ, SHHγ, and SHHδ) (Figure 6E) that generally follow age-specific patterns (Figure 6B). These SHH tumors show a clear age distribution with infant cases (<3 years) predominantly in SHHβ/γ, childhood cases (3-16 years) in SHHα, and adult cases (>17 years) in SHHδ (Figure 6C), with relatively balanced gender distribution across SHH subtypes (Figure 6D). Each subtype has different 5 year survival rates with SHHδ having the highest rates at around 80% followed by SHHβ, SHHγ, then SHHα with the worst at about 40%^14,36,37^ (Figure 6E).

Application of source topics showed several with high hierarchical association with SHH subgroup, specifically topics 2, 49, 22, and 46, with variable expression across other subgroups (Figure 7A). This gradient across subgroups suggests the granule cell precursor relationship with SHH medulloblastoma and varying degrees of developmental program retention across subgroups.

**Figure 7.**
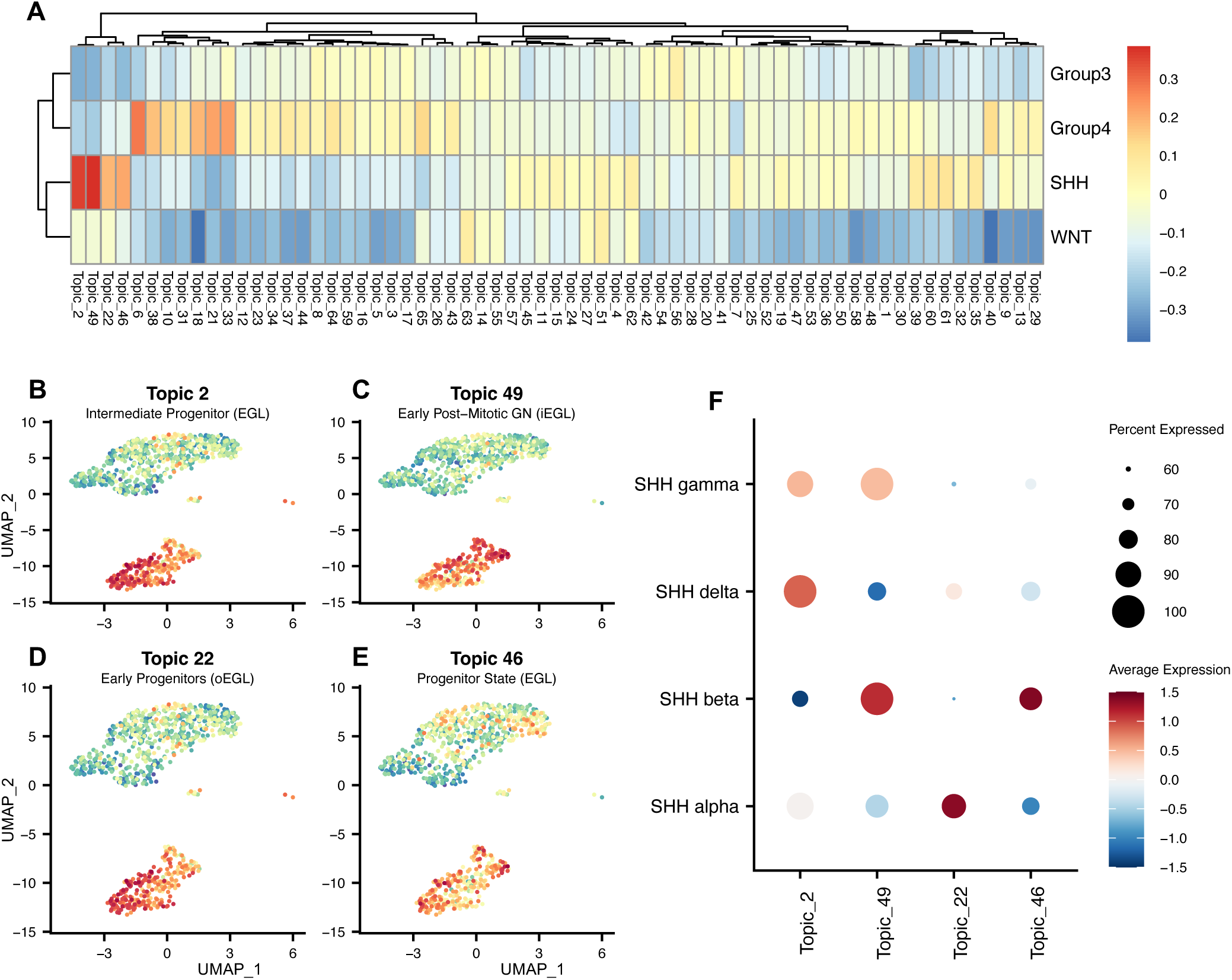
SHH subtypes express different developmental source topics and correspond to specific stages of EGL transitions. (A) Heatmap showing hierarchical clustering of 65 cerebellar developmental topics (columns) across medulloblastoma subgroups (rows). Color intensity represents scaled GSVA enrichment scores, with red indicating high expression and blue indicating low expression. (B-D) UMAP projections of topic expression scores across all 876 medulloblastoma samples, with color intensity representing GSVA enrichment scores. (B) Topic 2 (Intermediate Progenitor/EGL) (C) Topic 49 (Early Post-Mitotic GNs/iEGL) (D) Topic 22 (Early Progenitors/oEGL) (E) Topic 46 (Progenitor State/ EGL) (F) Dot plot quantifying topic association across SHH subtypes with dot size indicating percent of tumors under SHH subtype categorization with non-zero GSVA scaled score of the topics and color indicating average scaled GSVA score.

### SHH medulloblastoma subtypes associate with distinct developmental programs

The four SHH medulloblastoma subtypes showed distinct topic associations that corresponded with their age of onset. Each subtype appears to retain transciptional signatures from a specific stage of EGL development (associated with the topics), suggesting that the developmental timing at tumor initiation fundamentally shapes tumor biology.

SHHδ tumors occur predominantly in adults with TERT promoter mutations and showed preferential enrichment for Topic 2 (Figure 6B and E)^38^. This topic contains SHH pathway components *PTCH1* and *GLI2*, transcription factors *MEIS1*, *DACH1*, and *NFIA*, alongside additional markers whose functions are less characterized (*EGFEM1P*, *NAALADL2*, *XYLT1*) and are distinct for the adult tumors versus pediatric. SHHα tumors arise in children aged 3-16 years with frequent TP53 mutations and MYCN/GLI2 amplifications^39^. These tumors showed almost exclusive expression of Topic 22 (Figure 7C and E). This topic contained proliferation-associated genes *MKI67* (cell division marker), *CDK6* (cell cycle regulator), *EZH2* (chromatin remodeling), and *GLI2* (SHH signaling), alongside additional proliferation and cell division genes including *MPPED2*, *DIAPH3* (cytoskeleton regulation), *STARD13*, *KNTC1* (kinetochore function), and *CIT* (cytokinesis)^40^. This represents the most proliferative, outer EGL-like signature amont SHH subtypes which is consistent with this subtype’s aggressive clinical behavior and poor prognosis.

SHHβ tumors, arising in infants under 3 years with frequent SUFU mutations^39^, showed co-enrichment of Topic 46 (Figure 7E), containing neural developmental transcription factors *NFIA*, *NFIB*, *TCF4*, *CHD7*, and *MEIS1*. The presence of *ZEB1,* which regulates germinal zone exit, alongside markers of developmental plasticity suggests that these tumors retain characteristics of a progenitor state with sustained differentiation potential. These same SHHβ tumors also demonstrated predominant enrichment for Topic 49 (Figure 7C and E), which contained synaptic adhesion molecules (*LRRTM4*, *NLGN1*, *NLGN4X*), the *CSMD* family of cell surface glycoproteins (*CSMD1*, *CSMD2*, *CSMD3*), and additional neural development markers including *LHFPL3*, *NTNG1* (axon guidance), *AGBL4*, and *STK32B*. This co-enrichment pattern suggests SHHβ tumors arise from or retain characteristics of inner EGL cells initiating differentiation and synaptic development.SHHγ tumors, also occurring in infants, showed intermediate expression patterns with moderate enrichment for Topics 2 and 49, but lacking the strong Topic 46 co-enrichment observed in SHHβ (Figure 6E-F).

## DISCUSSION

### Topic modeling addresses fundamental challenges in comparative genomics

In the era of big data in genomics, several critical challenges continue to perplex the field especially in the context of statistical and machine learning: the lack of generalization of models to novel experimental conditions, the complexity of interpreting learned models (“black box”), and the application to complex cell dynamics. Here, we show how topic modeling can directly address these challenges and validate in the context of neurodevelopment and pediatric brain tumor origins.

The conservation of topics across datasets with different technologies, processing methods, and biological contexts as we show here with sci-RNA-seq3, SPLiT-seq, and bulk-RNA-seq, demonstrates that topic modeling captures fundamental biological information rather than dataset-specific artifacts. This generalization reflects the principle that cells coordinate gene expression in modules that drive specific biological functions^41,42^. Unlike approaches that require batch correction or data integration, this transferable topic modeling framework learns these modules from genetic signatures from source data and projects them onto target datasets through GSVA. This enables cross study comparison without computationally intensive integrations and dramatically reduces data loss as all genes are retained in the analysis.

In addition, the interpretability of topic modeling derives from its probabilistic, modular structure. Each topic consists of genes with quantified association strengths, enabling direct biological interpretation through pathway analysis, literature review, and spatial validation. Our EGL-associated topics map onto known developmental processes (proliferation (Topics 22, 2), glial specification (Topic 27), and neuronal differentiation (Topics 49, 61, 25, 46)) while revealing previously unrecognized transitional states within these broad categories. The topic-gene association values provide quantitative rankings that can be validated through orthogonal approaches such as in situ hybridization, as we demonstrated for nine marker genes in developing mouse cerebellum. This type of interpretability aligns with the goal of deriving biological insights from computational models rather than treating them as black boxes.

The ability to capture cell dynamics through transitional states represents perhaps the most significant advance that builds upon conventional clustering. Standard clustering fragments or entirely misses populations undergoing rapid state transitions due to their small size and intermediate expression profiles³². Topic modeling’s probabilistic assignments allow cells to express multiple topics simultaneously, naturally capturing the gradual nature of differentiation. Our identification of Topic 22 as a rhombic lip progenitor signature exemplifies this capability. These cells were present in multiple independent datasets but missed by conventional analysis due to their small numbers and transitional nature. The topic emerged not because we searched for rhombic lip markers, but because these genes consistently co-occurred in a subset of cells, revealing an underlying biological program invisible to discrete clustering approaches.

### Biological Insights into EGL Lineage Transitions and Specification

Our analysis reveals previously underappreciated complexity in the EGL while building upon conventional single cell and bulk RNA sequencing analysis. The identification of distinct transcriptional programs for the more proliferative progenitors (Topics 27 and 2) versus the oEGL-to-iEGL neuronal progression (Topics 22, 49, 61, 25) shows that these lineage decisions involve fundamentally different molecular mechanisms. The progenitor signatures appear more direct, with Topic 27 capturing multipotent glial progenitors. The presence of both *TNC* and *ETV5* (glial markers) alongside *SLC1A3* and *GDF10* (broader glial markers) in Topic 27 suggests these progenitors retain flexibility. The neuronal lineage shows complexity above conventional clustering with our topics capturing a developmental spectrum from proliferative oEGL through differentiating iEGL to migrating granule neurons. This progression involves at least five distinct transcriptional states (Topics 22, 49, 46, 61, 25). This reflects the nuances of granule neuron development, which requires massive clonal expansion in the oEGL, cell cycle exit and initial differentiation in the transition zone, further differentiation and synaptic specification in the iEGL, radial migration through multiple cerebellar layers, and finally synaptic integration with Purkinje cells and other neuronal populations. Each stage requires distinct transcriptional programs, which our topics successfully delineate.

### Developmental Timing and Tumor Biology

The subtype-specificity of topic expression within SHH medulloblastomas provides compelling evidence that developmental timing fundamentally shapes tumor transcriptional architecture. Rather than all SHH tumors arising from identical cellular origins with subsequent divergence, our results suggest that the developmental stage at tumor initiation, influenced by patient age, determines which transcriptional programs persist in the tumor itself.

Adult SHHδ tumors’ enrichment for Topic 2 suggests retention of an intermediate EGL progenitor state core SHH signaling components. These tumors arise in a cerebellum that has completed development, where any tumorigenic transformation must occur in rare residual or reactivated progenitors that existed during development. The combination of SHH pathway activity with mature transcription factor networks may provide alternative survival mechanisms when the primary SHH pathway is blocked, potentially explaining the variable response to therapeutic inhibitors observed clinically in adults^37,38^. In addition, the relatively uncharacterized markers *EGFEM1P*, *NAALADL2*, and *XYLT1* identified in this signature warrant further investigation as potential age-specific markers distinguishing adult from pediatric SHH tumors.

Infant SHHβ tumors’ enrichment for Topic 49 reflects their origin during active cerebellar development when the inner EGL is actively generating differentiating granule neurons engaged in synaptogenesis and migration. The abundance of synaptic adhesion molecules (*LRRTM4*, *NLGN1*, *NLGN4X*) and the complete *CSMD* family suggests these tumors arise from or retain characteristics of inner EGL cells that are locked in a specific developmental window focused on establishing neural connections. The co-enrichment of Topic 46’s neural transcription factors (*NFIA*, *NFIB*, *TCF4*, *CHD7*, *MEIS1*) indicates these tumors retain broader developmental plasticity, potentially contributing to their metastatic potential and these cells may maintain the migratory and adaptive programs of normal EGL development. The unique markers *LHFPL3*, *NTNG1*, *AGBL4*, and *STK32B* could serve as infant-specific biomarkers for diagnostic stratification.

Childhood SHHα tumors’ exclusive expression of Topic 22 represents a particularly aggressive phenotype where outer EGL-like proliferation programs dominate without accompanying differentiation signals. The concentration of cell cycle genes (*MKI67*, *CDK6*, *EZH2*) alongside factors associated with genomic instability (*KNTC1* overexpression linked to chromosomal instability, *DIAPH3* regulating cytoskeleton dynamics, *CIT* essential for cytokinesis) creates a molecular signature consistent with their dismal prognosis (5-year survival ∼41%)^14,43^. This hyperproliferative outer EGL-like state, combined with TP53 loss, generates uncontrolled proliferation and genomic instability that may explain why these tumors have the worst outcomes among SHH subtypes despite occurring in children who would otherwise have better regenerative capacity than adults. The dominance of proliferation markers suggests potential vulnerability to cell cycle inhibitors, particularly CDK4/6 inhibitors targeting *CDK6* or EZH2 inhibitors targeting the chromatin remodeling machinery^14,36–38^, though further in vitro validation followed by in vivo and clinical validation would be required.

SHHγ tumors’ intermediate expression pattern, with moderate enrichment for Topics 2 and 49 but lacking strong Topic 46 co-enrichment, may reflect their transitional position between infant and childhood tumor biology. This pattern suggests these tumors retain some developmental flexibility without being locked into either a more proliferative outer EGL-like state of SHHα or the inner EGL synaptic differentiation program of SHHβ. Further investigation with larger sample sizes and in vitro and in vivo validation is needed to fully characterize this subtype’s developmental relationships and clinical implications.

### Methodological Advances and Broader Applications

Beyond the specific biological insights, this work showcases topic modeling as a powerful tool for comparative and big-data genomics. The ability to transfer topics across datasets without batch correction, data integration, or shared processing pipelines solves challenges as the field generates ever-larger atlases with evolving technologies. This transferability corresponds to topics capturing genuine biological programs that are beyond the technical variations.

The framework’s ability to work across data modalities, as shown here in single cells to bulk tumors, is particularly valuable for translational research. Many clinical samples are only available as bulk RNA-seq due to historical technologies, sample quality, quantity, or cost constraints. Additionally, the current gold standard analysis for integrating single cell datasets is to reanalyze the original datasets with a consistent, current framework across the datasets. Original fastq files are not always available for multiple reasons and therefore not all prior datasets can be reanalyzed in this way. The topic analysis overcomes this limitation and provides a method for comparing and expanding the original analyses.

Our demonstration that developmental topics remain detectable and informative in bulk tumor samples opens new avenues for analyzing clinical cohorts and understanding how developmental programs contribute to diverse pathologies.

### Future Directions and Implications

The topics and statistical framework presented here have immediate applications beyond medulloblastoma and cerebellar development. Other pediatric brain tumors suspected to have developmental origins, including other embryonal tumors, pediatric gliomas, and ependymomas, could be analyzed using the same cerebellar developmental topics or topics derived from appropriate developmental atlases. Adult brain tumors might also retain or reactivate developmental programs detectable through this approach.

More broadly, this framework could transform how we approach the relationship between development and disease across all organ systems. As developmental atlases become available for heart, lung, kidney, and other organs, the same methodology could identify which developmental programs persist in congenital anomalies, contribute to adult disease, or become reactivated in cancer. The key insight is that development provides a template for understanding disease and by comprehensively mapping normal developmental programs we continue to create a reference frame for understanding pathology.

Additional future directions includes further in vitro validation of the genetic signatures within the topics explored here to further strengthen the applicability of these programs. In this case, exploration of the top 5-10 genes from the topics associated with the RL and bifurcating trajectories, particularly those distinguishing oEGL from iEGL states, can be explored in new human developmental tissue, mouse models, and medulloblastoma samples. Patient-derived organoid models could be particularly valuable for testing whether manipulation of stage-specific EGL programs alters tumor behavior or drug sensitivity.

As single-cell technologies generate ever-larger datasets, approaches that extract meaningful patterns without computationally intensive integrations become necessary. Topic modeling meets this need by providing a scalable framework for building cumulative knowledge from the exponentially growing corpus of transcriptomic data. The topics and methods presented here are immediately applicable to other biological systems and malignancies, offering a path forward for integrating datasets across different biological contexts and technologies. By using this framework in the context of neurodevelopment and early cancer pathogenesis, we showcase how hard to identify transitional cellular populations can be characterized efficiently. This characterization was transferred to tumor data and we showed persisting developmental features from specific stages of EGL development that may drive medulloblastoma subtype diversity and could inform personalized treatment strategies.

### Limitations of the study

Limitations of this study include that topic number selection requires balancing granularity with interpretability. Different values might reveal additional insights. The uncharacterized markers and marker combinations found with these topics require further functional validation to confirm their roles in development and disease in human. Our analysis focused on a limited pre-natal development time whereas earlier and later stages would complete the developmental picture. Despite these limitations, the framework showcased valuable insights including conserved programs across independent datasets, cell populations hidden from commonly used clustering approaches, and connected developmental programs to cancer subtypes

## RESOURCE AVAILABILITY

### Lead contact

Further information and requests for resources and reagents should be directed to and will be fulfilled by the lead contact, Siobhan Pattwell.

### Materials availability

#### Data and code availability

All original code for topic modeling is deposited at https://doi.org/10.5281/zenodo.17274488. Publicly available datasets analyzed in this study include: Aldinger et al. human fetal cerebellar dataset (GEO: GSE162170), Cao et al. human fetal atlas (GEO: GSE156793), and Arora et al. medulloblastoma bulk RNA-seq (https://doi.org/10.1101/2024.10.21.619495). Mouse cerebellar single-cell data was obtained from Qiu et al. (GEO: GSE186069). In situ hybridization images were accessed from the Allen Developing Mouse Brain Atlas (https://developingmouse.brain-map.org/).

## ACKNOWLEDGMENTS

This work was supported in part by the National Institutes of Health, National Library of Medicine (NLM) University of Washington Biomedical Informatics and Data Science Research Training Program (Grant Nr LM 007442) and (Grant Nr 5K22CA258953 through SSP). KJM is supported by the National Institutes of Health R01CA270785 and R37NS095733. PH is supported by the National Institutes of Health R21NS133390, R21NS138661 and R21NS142543 The content is solely the responsibility of the authors and does not necessarily represent the official views of the National Institutes of Health. We thank the Allen Institute for Brain Science for providing access to the Developing Mouse Brain Atlas data.

## AUTHOR CONTRIBUTIONS

Conceptualization, A.R. and S.S.P.; methodology, A.R., S.S.P., J.G., K.G., and S.S, PH, and KJM.; Investigation, A.R., S.S.P., J.G., K.A.A, K.J.M, P.H..; writing--original draft, A.R. and K.G., and S.S.; writing- -review & editing, S.S.P, J.G, K.A.A., K.J.M, P.H, K.G., S.S., J.S., D.J., L.M.G., S.A.; funding acquisition, S.S.P., A.R.; resources, S.S.P..; supervision, S.S.P, J.G.

## DECLARATION OF GENERATIVE AI AND AI-ASSISTED TECHNOLOGIES

No Generative AI or AI-assisted technologies were used in the construction of this paper or study.

## STAR★METHODS

### KEY RESOURCES TABLE

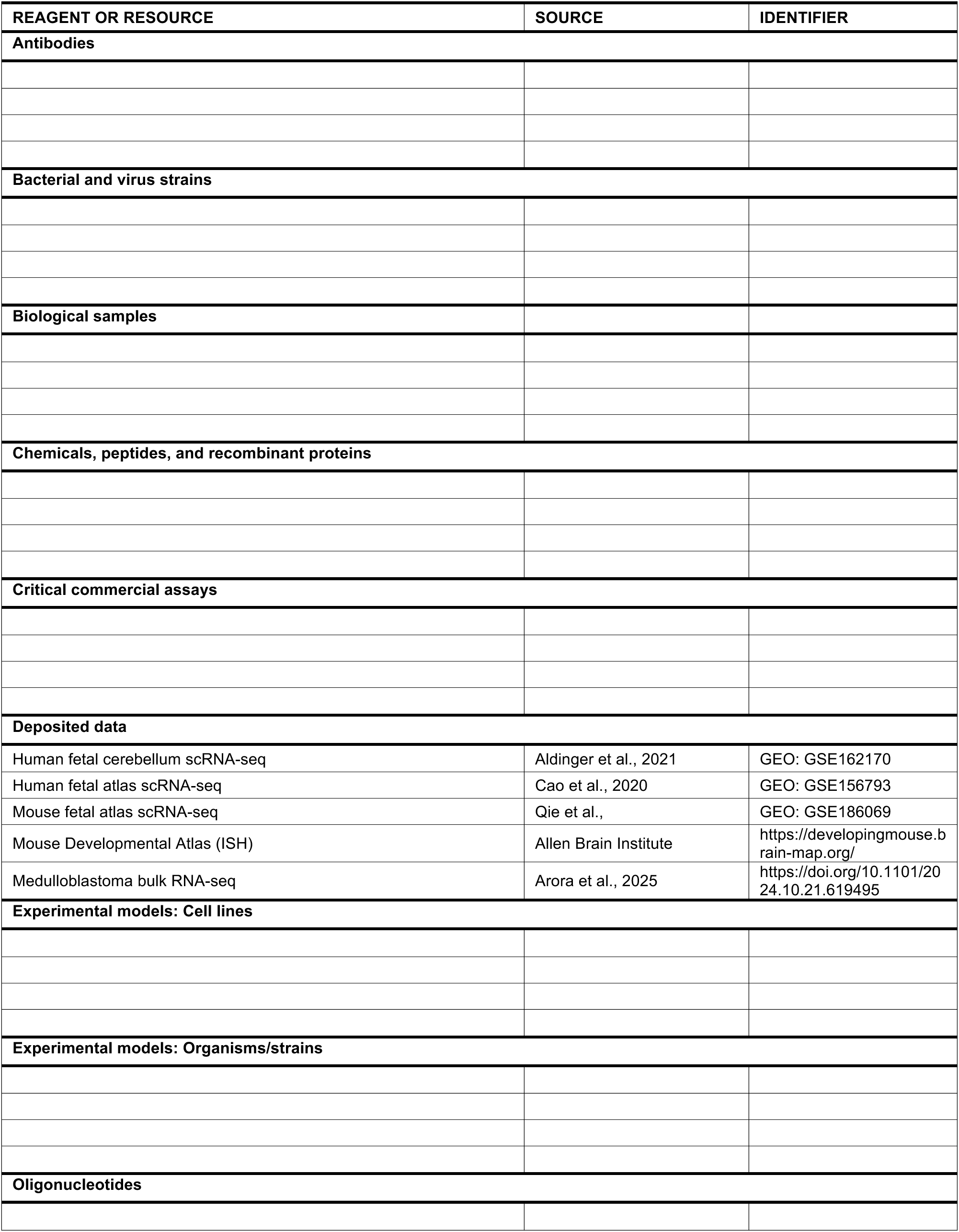
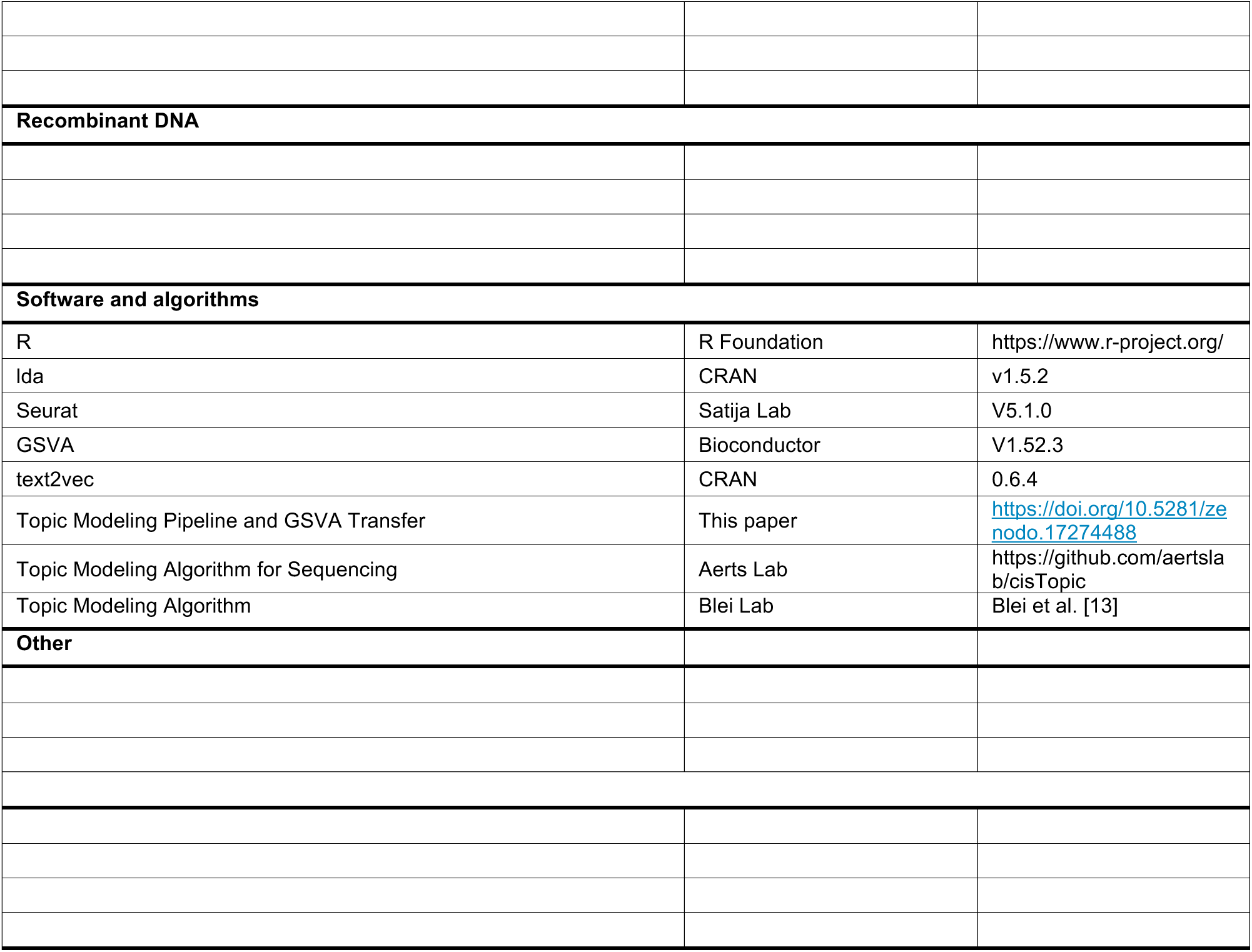

### EXPERIMENTAL MODEL AND STUDY PARTICIPANT DETAILS

#### Human Fetal Tissue Datasets

The source dataset consisted of human fetal cerebellar tissue from 13 specimens spanning post-conception weeks (PCW) 9-21, obtained from the Aldinger et al. study (2021). This dataset was subset to include timepoint PCW 17 with 15,556 nuclei processed using SPLiT-seq technology. The validation dataset consisted of human fetal cerebellar cells from 5 specimens spanning PCW 12-17, obtained from the Cao et al. human cell atlas (2020). This dataset included 119,954 cerebellar nuclei from PCW 12.5 identified from the larger atlas, processed using sci-RNA-seq3 technology.

#### Medulloblastoma Samples

Bulk RNA sequencing data from 876 medulloblastoma tumors were obtained from the updated reference landscape of human brain diseases (Arora et al., 2023). Samples included all four molecular subgroups: WNT (n=70), SHH (n=274), Group 3 (n=144), and Group 4 (n=388). SHH subgroup samples were further stratified into four subtypes (α, β, γ, δ) based on age and molecular features.

### METHOD DETAILS

#### Data Acquisition and Preprocessing—Single Cell Data

Single-cell RNA sequencing data were downloaded from Gene Expression Omnibus (GEO). The Aldinger et al. dataset was shared by the original authors and Cao et al. dataset (GSE156793) were loaded as Seurat objects. Mouse single cell data was downloaded from (GSE186069) from Qiu et al. and randomly downsampled to 100,000 cells for computational efficiency. When normalization was enabled, data were normalized using Centered Log Ratio (CLR) transformation via Seurat’s NormalizeData function. Variable features were identified using FindVariableFeatures, with the number of features set to total genes in the RNA assay (18,676 genes for Aldinger dataset; 30,000 for Cao dataset; 19,352 for Qiu dataset).

#### Data Acquisition and Preprocessing—in situ hybridization

Mouse brain in situ hybridization images were obtained from the Allen Developing Mouse Brain Atlas (https://developingmouse.brain-map.org/). Expression patterns of selected topic marker genes were examined at postnatal day 4 (P4) in sagittal sections. Genes examined included PAX6, BOC, RELN, MEIS1, EPHA6, GRIK2, NFIA, TCF4, and ERBB4, representing markers across the EGL developmental topics available through the atlas.

#### Topic Modeling Implementation - Cross-Species Gene Name Alignment

For transferring human source topics to mouse dataset, gene name alignment was performed by converting mouse genes to human orthologs using gProfiler (https://biit.cs.ut.ee/gprofiler/). Only genes with confident one-to-one ortholog mappings were retained for analysis. The conversion table is provided as Supplementary Table S4.

#### Topic Modeling Implementation--Matrix Preparation

The sparse expression matrix was extracted from the RNA assay. To prepare data for LDA analysis, the sparse matrix was multiplied by 10 and rounded to convert continuous expression values to discrete count data suitable for the collapsed Gibbs sampler. Data were reformatted into cell-specific lists containing gene indices and expression values as required input for lda.collapsed.gibbs.sampler.

#### Iterative LDA Training

LDA models were trained using the collapsed Gibbs sampling algorithm implemented in the lda package (v1.5.2) with the following parameters: Alpha (document-topic concentration): 50, Beta/Eta (topic-word concentration): 0.1, Number of iterations: 100, Burn-in period: 300 iterations, Random seed for reproducibility. Models were trained across a range of topic numbers (K=20 to K=100, incrementing by 5) to enable optimal model selection. Each model was saved as an RDS file for subsequent analysis.

#### Topic Gene Set Extraction

Topic-specific gene signatures were extracted using lda::top.topic.words function. Genes were ranked by probability scores (by.score = TRUE) within each topic. The top 50 genes per topic were extracted and formatted as gene sets named “Sample_Topic_N” (e.g., “Ald17_Topic_22”) to maintain sample and topic identity.

#### Cross-Dataset Topic Transfer via GSVA

For transferring human source topics to mouse dataset, gene name alignment was performed by converting mouse genes to human using gProfiler^44^ (Supplementary table S4). Gene Set Variation Analysis (GSVA v1.52.3) was employed to calculate single-sample gene set enrichment scores, enabling topic activity quantification at individual cell level. Topic gene sets were scored against log2-transformed expression data using gsvaParam function with maximum difference method (maxDiff=TRUE). This non-parametric approach calculates enrichment by computing maximum deviation of empirical cumulative distribution function of gene set genes from remaining genes. GSVA scores represent relative enrichment of each topic’s gene signature in individual cells or bulk samples, with positive scores indicating topic activation. Scores were added to dataset metadata, enabling cross-sample comparison and identification of conserved biological programs across independent datasets.

### QUANTIFICATION AND STATISTICAL ANALYSIS

#### Model Selection via Perplexity Analysis

Optimal topic number was determined using perplexity. For each model, document-term matrices were reconstructed and topic-word distributions were extracted and L1-normalized. Document-topic distributions were derived from model document sums and similarly normalized. Perplexity was calculated using the text2vec::perplexity function. The optimal K=65 topics was selected where perplexity began to plateau, balancing model complexity with interpretability.

#### Statistical Analysis

All statistical analyses were performed in R (v4.4.0). Topic enrichment scores across cell types were visualized using dot plots showing average expression and percentage of cells expressing each topic. Hierarchical clustering was performed using Ward’s method on Euclidean distances. Heatmaps were generated using pheatmap with row and column clustering.

#### Data Visualization

UMAP projections were generated using Seurat’s RunUMAP function with default parameters. Topic expression overlays were created by mapping GSVA scores to UMAP coordinates. All other plots were generated using ggplot2.

## SUPPLEMENTAL INFORMATION

**Figure S1.**
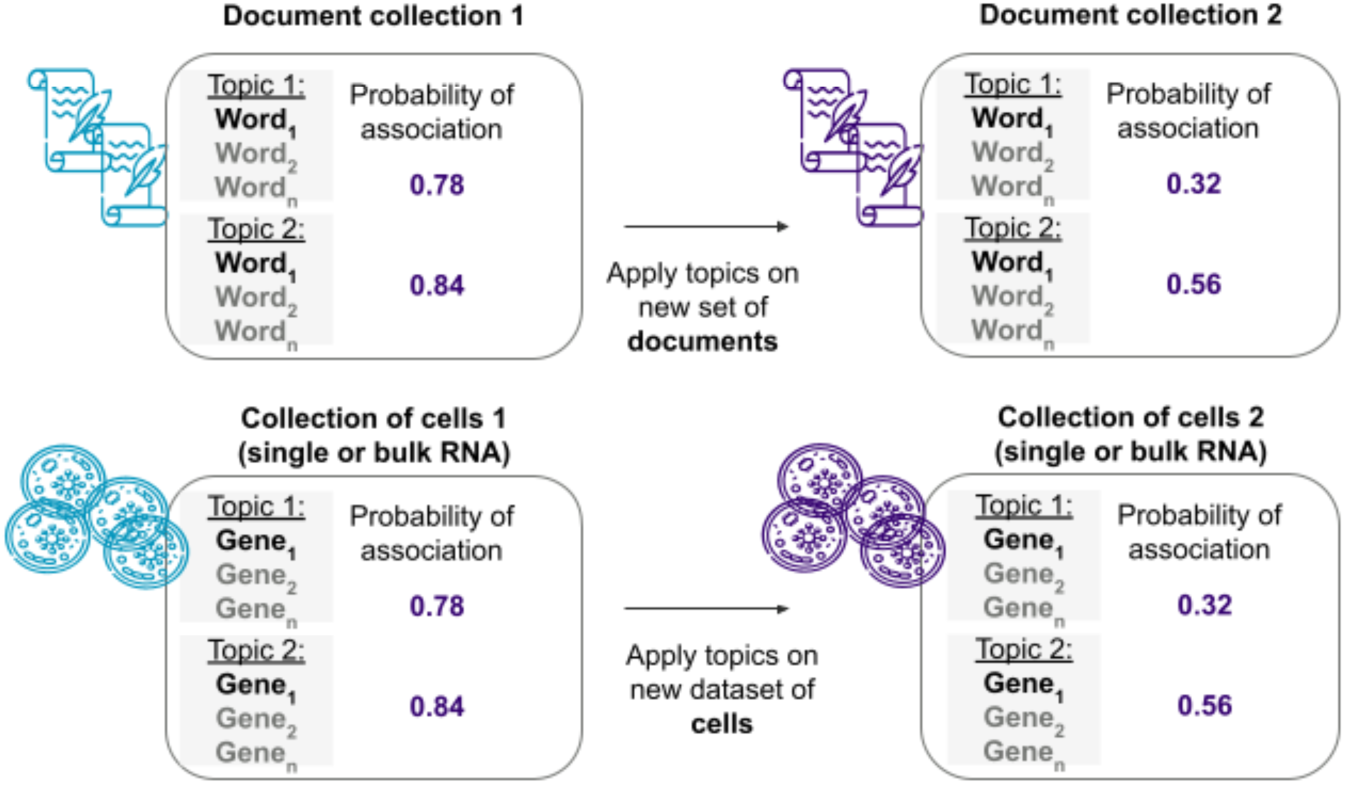
Analogy between topic modeling in document analysis and gene expression analysis in single-cell or bulk RNA sequencing. Top row: Conventional topic modeling identifies topics (collections of co-occurring words) across document collections. Bottom row: The same mathematical framework can be applied to identify gene expression programs (collections of co-expressed genes) across cell populations. Topics discovered in documents map to gene expression patterns in cells, with associated probability values indicating the strength of each topic/pattern.

**Figure S2.**
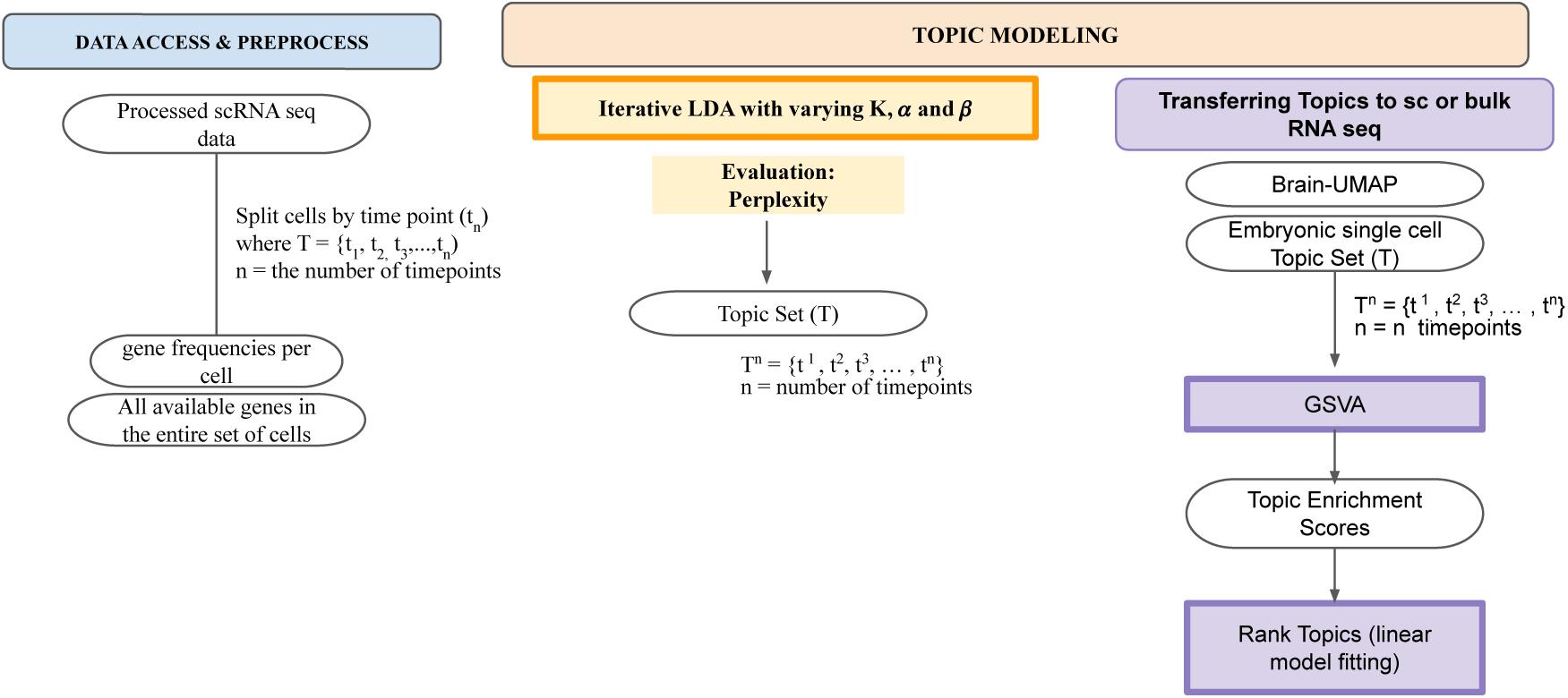
Workflow for applying topic modeling to single-cell and bulk RNA sequencing data. Left panel: Data preprocessing pipeline showing cell stratification by timepoint and gene frequency calculation. Right panel: Topic modeling pipeline using iterative Latent Dirichlet Allocation (LDA) with parameter optimization (K, α, β), followed by topic transfer to either Brain-UMAP or embryonic single-cell datasets. The workflow concludes with Gene Set Variation Analysis (GSVA), topic enrichment scoring, and ranking via linear model fitting to identify biologically relevant gene expression programs.

**Figure S3.**
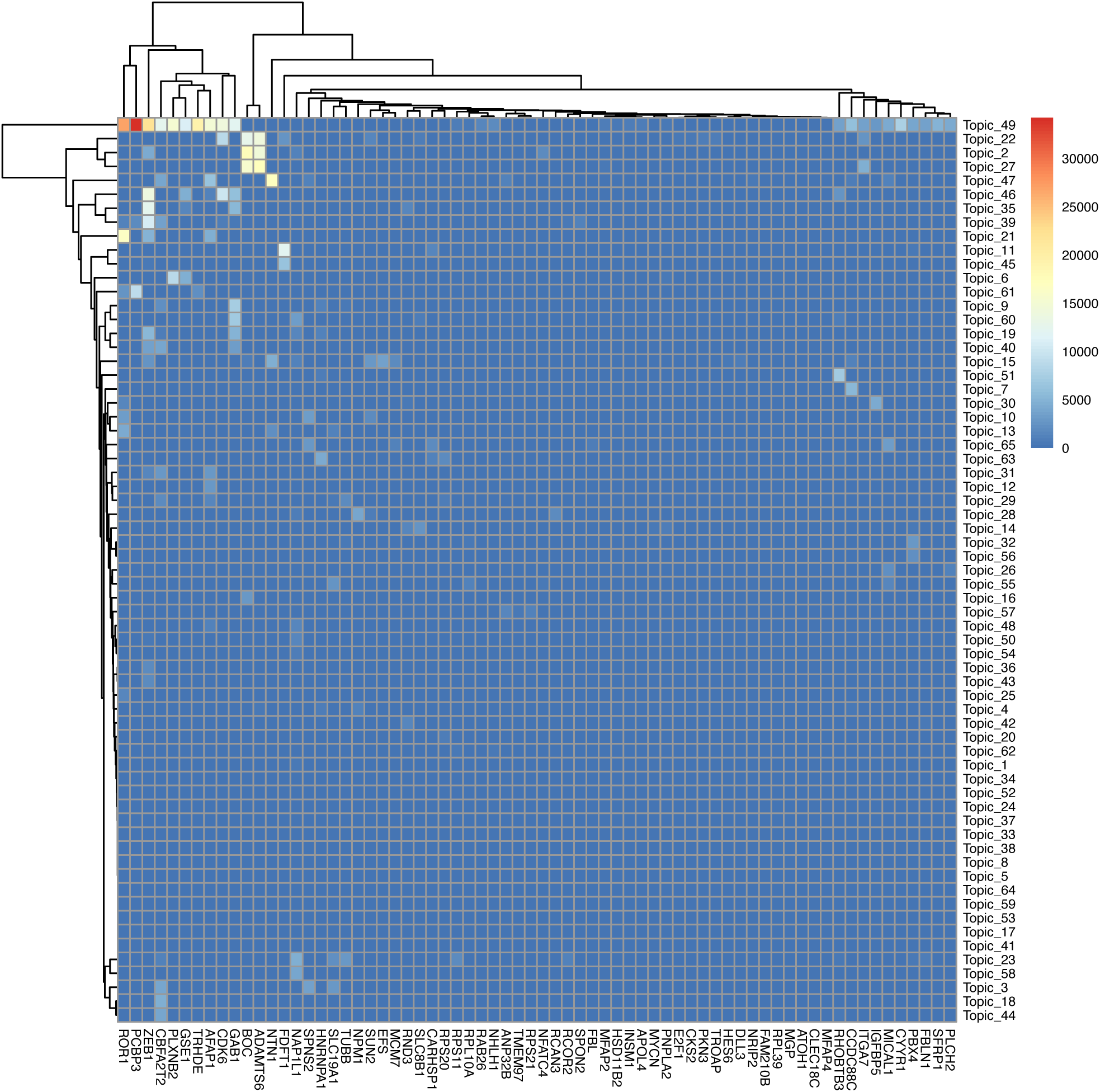
Heatmap of gene-topic association values for the source topics for the top differentially expressed genes (DEGs) from the external granular layer (EGL) from Aldinger et al. Genes (columns) represent the top EGL DEGs identified in the Aldinger et al.^1^ study on spatial and cell type transcriptional landscape of human cerebellar development. Rows represent topics derived from LDA modeling of the same Aldinger et al. dataset. Color intensity indicates the strength of gene-topic associations. Hierarchical clustering reveals that EGL marker genes show selective enrichment in specific topics, suggesting these topics specifically in topic 49.

**Figure S4.**
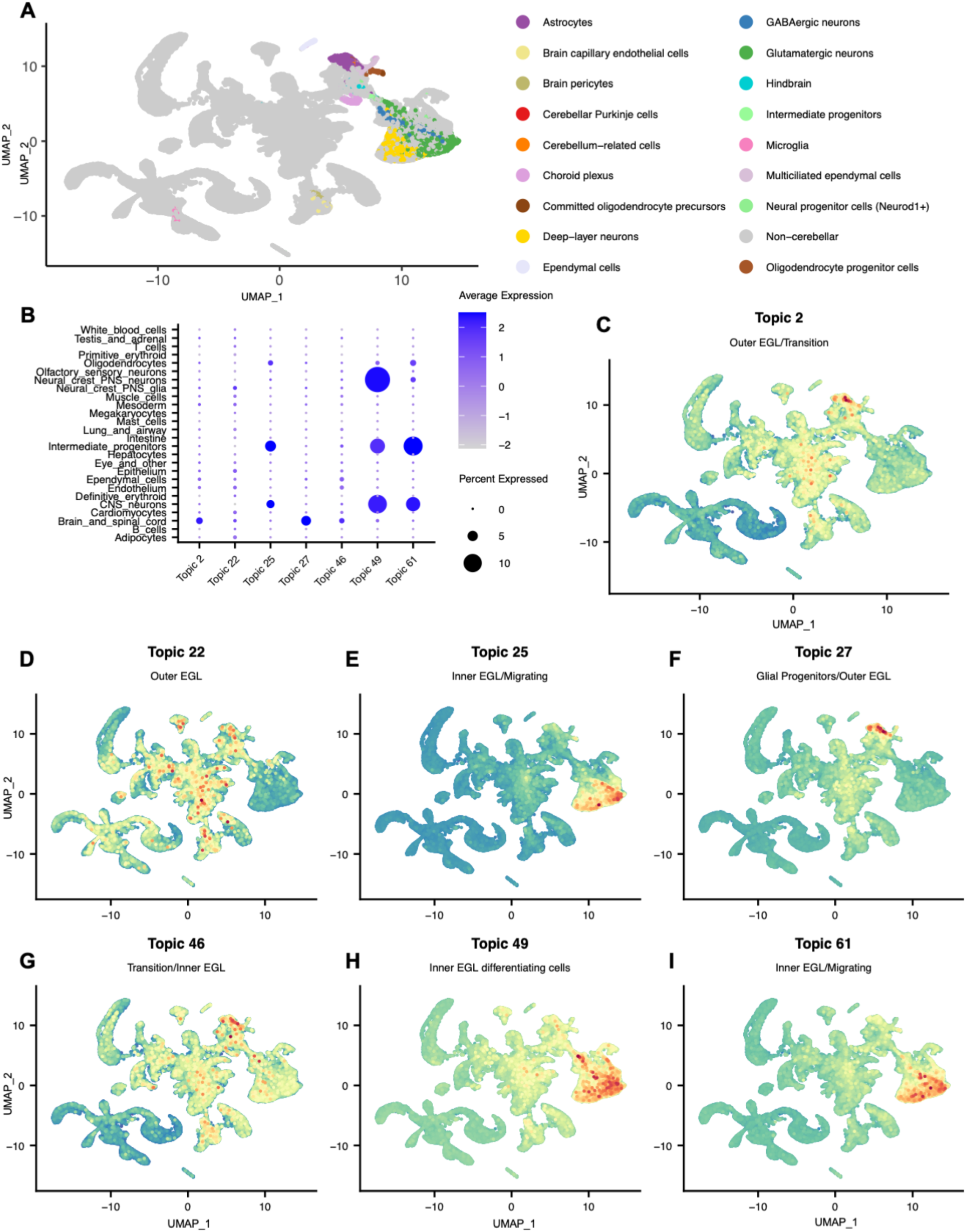
Cross species validation of source topics (human cerebellar developmental topics) transferred to single cell developmental mouse atlas using GSVA. (A) UMAP visualization of 100,000 mouse cerebellar cells (downsampled from Qiu et al. dataset) colored by cell type annotations. Human gene symbols were converted to mouse orthologs using gProfiler for cross-species topic transfer. (B-I) UMAP projections showing GSVA enrichment scores for transferred human topics in mouse cerebellum: (B) Topic 27 (Early Glial Progenitors) (C) Topic 2 (Intermediate Progenitor) (D) Topic 22 (Early Progenitors/oEGL) (E) Topic 49 (Early Post-Mitotic GNs) (F) Topic 61 (Post-Mitotic Maturing GNs) (G) Topic 25 (Transitioning GN) (H) Topic 46 (Progenitor State)

